# Age and obesity-driven changes in the extracellular matrix of the primary tumor and metastatic site influence tumor invasion and metastatic outgrowth

**DOI:** 10.1101/2023.08.24.554492

**Authors:** Sydney J. Conner, Justinne R. Guarin, Hannah B. Borges, Kenneth J. Salhany, Diamond N. Mensah, Grace A. Hamilton, Giang H. Le, Madeleine J. Oudin

## Abstract

Younger age and obesity increase the incidence and metastasis of triple-negative breast cancer (TNBC), an aggressive subtype of breast cancer. The extracellular matrix (ECM) promotes tumor invasion and metastasis. We characterized the effect of age and obesity on the ECM of mammary fat pads, lungs, and liver using a diet-induced obesity (DIO) model. At 4 week intervals, we either injected the mammary fat pads with allograft tumor cells to characterize tumor growth and metastasis or isolated the mammary fat pads and livers to characterize the ECM. Age had no effect on tumor growth but increased lung and liver metastasis after 16 weeks. Obesity increased tumor growth starting at 12 weeks, increased liver metastasis only at 4 weeks, and weight gain correlated to increased lung but not liver metastasis. Utilizing whole decellularized ECM coupled with proteomics, we found that early stages of obesity were sufficient to induce changes in the ECM composition and invasive potential of mammary fat pads with increased abundance of pro-invasive ECM proteins Collagen IV and Collagen VI. We identified cells of stromal vascular fraction and adipose stem and progenitor cells as primarily responsible for secreting Collagen IV and VI, not adipocytes. We characterized the changes in ECM in the lungs and liver, and determined that older age decreases the metastatic potential of lung and liver ECM while later-stage obesity increases the metastatic potential. These data implicate ECM changes in the primary tumor and metastatic microenvironment as mechanisms by which age and obesity contribute to breast cancer progression.

**Significance:** Younger age and obesity increase the incidence and metastasis of triple-negative breast cancer. Our data suggest that changes in the breast, lung and liver ECM are driving some of these effects.

## Introduction

Obesity is a growing pandemic. The prevalence of obesity in adults in the United States has been increasing by approximately 2% per year since 2010 [1]. Now, more than 3 in 5 women are overweight or obese, with nearly 12% having severe obesity [1]. In women, obesity is a risk factor for a higher incidence of breast cancer and an overall poorer prognosis [2], [3]. Obesity is linked to a decreased overall breast cancer survival rate [3]–[5] due to higher rates of metastasis [3], [6], and resistance to chemotherapy [6]–[8], a standard treatment for breast cancer. One mechanism of obesity-induced breast cancer progression is through elevated estrogen secretion by adipose tissue, specifically in obese postmenopausal patients [9], promoting hormone receptor-driven progression [10]. However, obesity has also been linked to increased incidence and progression of triple-negative breast cancer (TNBC), which lacks expression of progesterone, estrogen and HER2 receptors[6], [7], [11], [12]. TNBC is an aggressive subtype of breast cancer characterized by a high risk of metastasis, a lack of targetable receptors which makes it difficult to treat, and subsequently a poor survival rate 10% lower than non-TNBC [6], [11]–[13]. Age, which is associated with an increase in abdominal obesity [14], is also a risk factor for an increased incidence of breast cancer, with women over 60 having the highest incidence as well as poor prognosis [15]. However, this is not the case for TNBC, which is the most commonly diagnosed subtype in women under 40 [16]. Thus, there is a need to understand how obesity and age contribute to TNBC progression and metastasis to better treat this aggressive subtype in the growing obese population.

Obesity is a systemic disease that causes chronic low-grade inflammation and dysregulation throughout the body [17], [18]. The breast is comprised primarily of adipose tissue, which contains various cell types including, epithelial cells, immune cells, fibroblasts, adipose stromal cells, adipose stem and progenitor cells (ASPCs), and adipocytes. These cells are embedded in a meshwork of extracellular matrix (ECM) proteins that provide physical support and biochemical signals that influence cell behavior. Obesity causes physiological changes in breast adipose tissue function and ECM composition due to inflammation and adipocyte dysregulation [17]. During weight gain, adipose tissue expands through adipocyte hyperplasia (generation of new adipocytes) and adipocyte hypertrophy (enlarging of adipocytes). This limits the diffusion of oxygen and nutrients to distant parts of the tissue causing inflammation, immune cell infiltration, ECM deposition, and ultimately fibrosis [19]. Many of these changes have been shown to contribute to increased cancer incidence and progression. We and others have investigated the role of the ECM in breast cancer progression, which is well known to promote local invasion and metastasis [20]. Specifically, obesity-dependent changes in the mammary fat pad ECM have been shown to promote tumor cell invasion [21], [22]. Seo et al. found that ECM secreted by adipose stromal cells isolated from mammary fat pads of the ob/ob obese genetic mouse model compared to lean mice had an increased abundance of fibronectin and collagen cross-linking, leading to increased breast cancer cell invasion via mechanosignaling [21]. We previously demonstrated that whole decellularized mammary fat pad ECM, isolated from obese mice fed a high-fat diet (HFD) for 16 weeks compared to lean chow diet (CD) fed mice, drives tumor cell invasion. Additionally, we found the ECM protein Collagen VI as more abundant in obese mammary glands and as a driver of breast cancer invasion [23]. The cells responsible for changes in ECM production in the obese mammary gland are not well known. In addition, the role of obesity-dependent changes in the ECM at metastatic sites is not well understood. The two most common sites of breast cancer metastasis are the lungs and liver [24]. In murine models of diet-induced obesity (DIO) and breast cancer, HFD has been shown to promote primary tumor growth [25]–[30], lung metastasis [25], [26], [29], [31], and liver metastasis [31]. Obesity is known to affect the liver through lipid accumulation in hepatocytes called steatosis [18], [32], although the association between liver steatosis and breast cancer metastasis in humans is not clear [33], [34].

Further, the relationship between weight gain, ECM changes, and breast cancer metastasis in murine DIO is not well understood. It has been shown that in a murine DIO model after 4 weeks of HFD, there is a significant increase in body weight, hyperglycemia, and adipocyte hypertrophy [18], [35]. After 8 weeks, there is an increase in insulin levels [32] and immune cell infiltration into the adipose tissue [18], [32], and at 12 weeks, lipid accumulation begins in the liver [18], [32]. Thus, obesity-dependent changes are progressing even before the mouse is characterized as obese at 16 weeks, yet few studies have investigated how earlier stages of obesity (prior to 8 weeks on HFD) promote breast cancer progression or when obesity-dependent changes in the ECM that drive breast cancer invasion and metastasis occur. In humans, obesity also occurs on a spectrum; therefore, understanding how different stages of weight gain impact tumor formation and metastasis is important. Lastly, the majority of breast cancer and obesity studies use ovariectomized female mice to mimic the postmenopausal state, but TNBC is commonly diagnosed in women under 40 (premenopausal women). In addition to menopausal status, age has also been shown to have a greater impact on the expression of ECM and tissue remodeling genes in adipose stromal vascular cells than even obesity in mice [32] and been associated with priming the metastatic niche [36]. Thus, determining the effects of age and obesity on the ECM and tumor progression in murine models is critical.

In this study, we sought to characterize the effect of age and obesity on local breast adipose tissue and at metastatic sites to determine how these progressive changes in the microenvironment promote TNBC invasion and metastasis. Age had a significant effect on metastasis to the lungs and liver, but not tumor growth. Obesity-induced significant tumor growth and metastasis to the lungs, with effects on liver metastasis only seen early on. Early stages of obesity were sufficient to induce distinct differences in the ECM composition of mammary fat pads that are associated with increased tumor cell invasion. The ECM proteins Collagen IV and Collagen VI, previously implicated in driving TNBC invasion, were enriched with age and obesity. We found that both Collagen IV and VI are primarily secreted by ASPCs from adipose tissue and not adipocytes. Lastly, we found that longer time on HFD was required to induce pro-invasive changes in the lung and liver ECM. Overall, our studies show that both age and diet can impact ECM-driven effects on tumor progression and metastasis in TNBC.

## Results

### Age and high-fat diet affect tumor growth and metastasis to the lungs and liver

To investigate how age and obesity promote TNBC progression, we injected a murine adenocarcinoma cell line (E0771) into the 4^th^ left mammary fat pad of 10-week-old female C57BL/6 mice who had been fed either HFD or age-matched control CD for 4, 8, 12, and 16 weeks. Following tumor initiation, mice were kept on diet, weighed and tumors were measured weekly for 5 weeks (Fig 1A). Using non-tumor bearing mice, we confirmed that total body weight, adipocyte size, and fat pad weight were significantly increased on HFD starting at 4 weeks (Fig 1B, S1A-C). We also observed that our CD mice were gaining significantly more body weight as they aged (Fig S1D), but there was no significant difference in adipocyte size or fat pad weight (Fig S1A-C). Tumors grew faster and were significantly larger when mice had been fed HFD for 12 and 16 weeks prior to initiation of the tumor compared to age-matched CD mice (Fig 1C, D). Due to the heterogeneity of weight gain in both the CD and HFD groups, we examined whether there was a correlation between weight, uncoupled from duration or type of diet, and tumor volume. There was a positive correlation between the weight of mice when tumors were initiated and tumor volume at the terminal endpoint (Fig 1E). These data show that age itself did not affect tumor growth. While both age and HFD affected total body weight, only HFD for at least 12 weeks promoted primary tumor growth.

**Figure 1.**
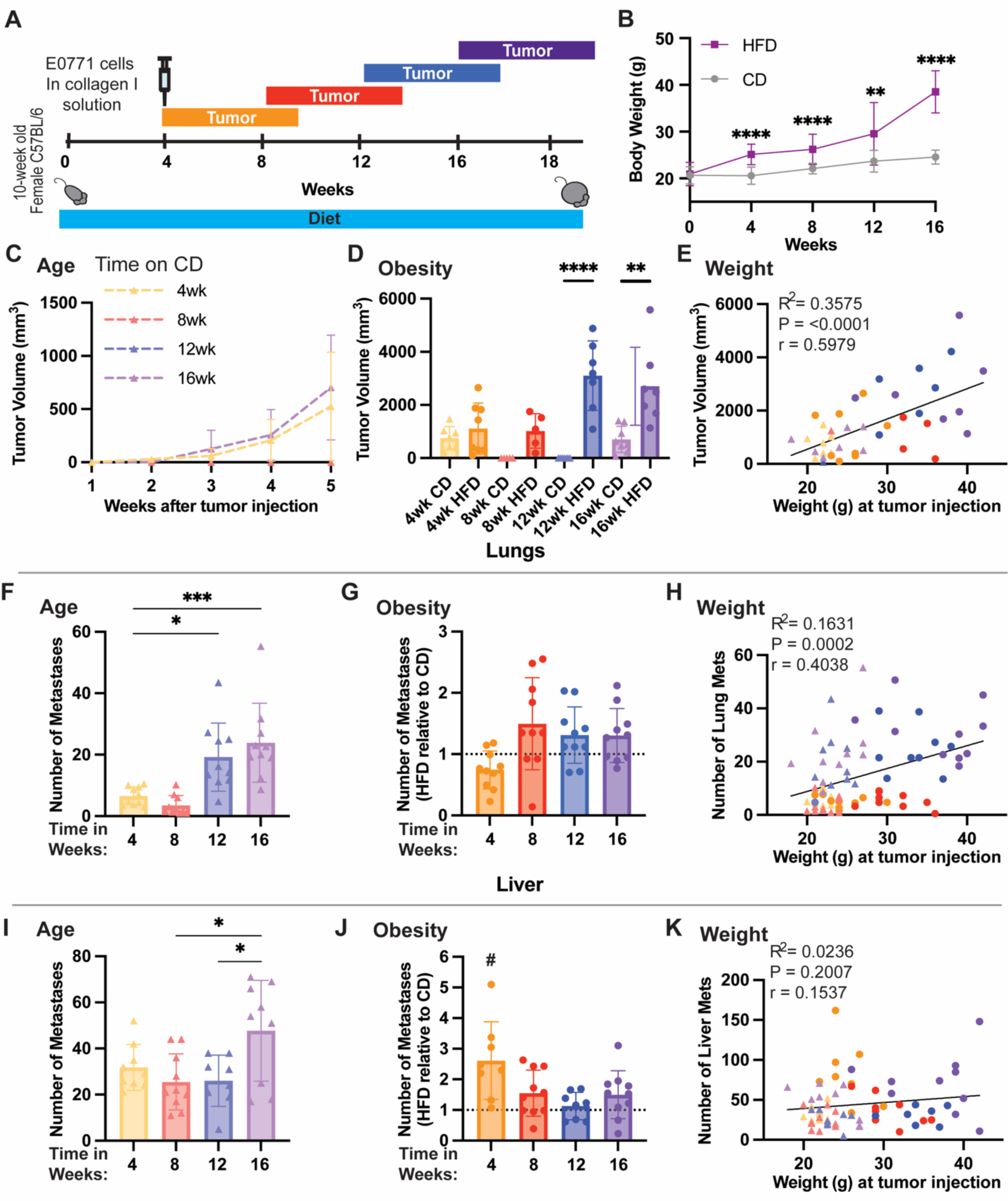
HFD-induced tumor growth. **(A)** Schematic depicting experimental paradigm where female mice were fed CD or HFD for 4, 8, 12, or 16 weeks, then injected with E0771 cells, and tumors were grown for 5 weeks. **(B)** Body weight of mice over time fed CD or HFD. **(C)** Tumor volume over time of tumors injected in mice fed CD for 4, 8, 12, and 16 weeks. **(D)** Tumor volume at endpoint 5 weeks after tumors were injected in mice fed HFD or CD for 4, 8, 12, and 16 weeks (P**<0.01, P***<0.001, P****<0.0001). **(E)** Correlation between tumor volume and mouse body weight at the time of tumor injection. **(F)** Number of lung metastases at endpoint 5 weeks after tumors were injected in mice fed CD for 4, 8, 12, and 16 weeks **(G)** Number of lung metastases on HFD relative to CD **(H)** Correlation between number of lung metastases (mets) at the endpoint and mouse body weight at the time of tumor injection. **(I)** Number of liver mets at endpoint 5 weeks after tumors were injected in mice fed CD for 4, 8, 12, and 16 weeks **(J)** Number of liver mets on HFD relative to CD **(K)** Correlation between the number of liver mets at the endpoint and mouse body weight at the time of tumor injection. Significance was determined by Ordinary one-way ANOVA with Tukey multiple comparisons to compare between CD groups (P*<0.05, P**<0.01, P***<0.001, P****<0.0001). An unpaired t-test was used to compare age-matched CD and HFD groups (#P<0.05). Data is shown as mean with SD n = 8-10 animals/group. Significance was determined using an unpaired t-test to compare age-matched CD and HFD groups.

We next investigated the effect of age and obesity on metastasis in two of the most common sites of TNBC metastasis: the lungs and liver. There were more metastases in the lungs when tumor cells were injected 12 and 16 weeks after starting CD than at 4 and 8 weeks (Fig 1F, S2A), suggesting that age increases lung metastasis in healthy mice. To account for these age-related differences in lung metastasis when comparing HFD groups, we quantified the number of metastases to the lungs on HFD relative to CD for each time point. HFD did not significantly increase lung metastasis compared to CD at any given time point (Fig 1G). We then wanted to see if weight, uncoupled from duration or type of diet, correlated with lung metastasis. Our data showed a significant positive correlation between the weight of mice when tumors were initiated and the number of lung metastases (Fig 1H). In the liver, there was a significant increase in metastasis between 8 and 16 weeks and 12 and 16 weeks on CD (Fig 1I), suggesting that age can increase liver metastasis, but only after 16 weeks. To dissect the effects of diet on each time point, we quantified the number of liver metastases on HFD relative to age-matched CD. In the liver, there was a significant increase in the number of metastases on HFD compared to CD but only at 4 weeks on HFD (Fig 1J, S2B). There was no correlation between weight at tumor initiation and the number of metastases in the liver (Fig 2K). These data indicate that in the lungs, age had a more pronounced effect on metastasis than HFD and in the liver early stages of HFD had the same pro-metastatic effect as older age.

**Figure 2.**
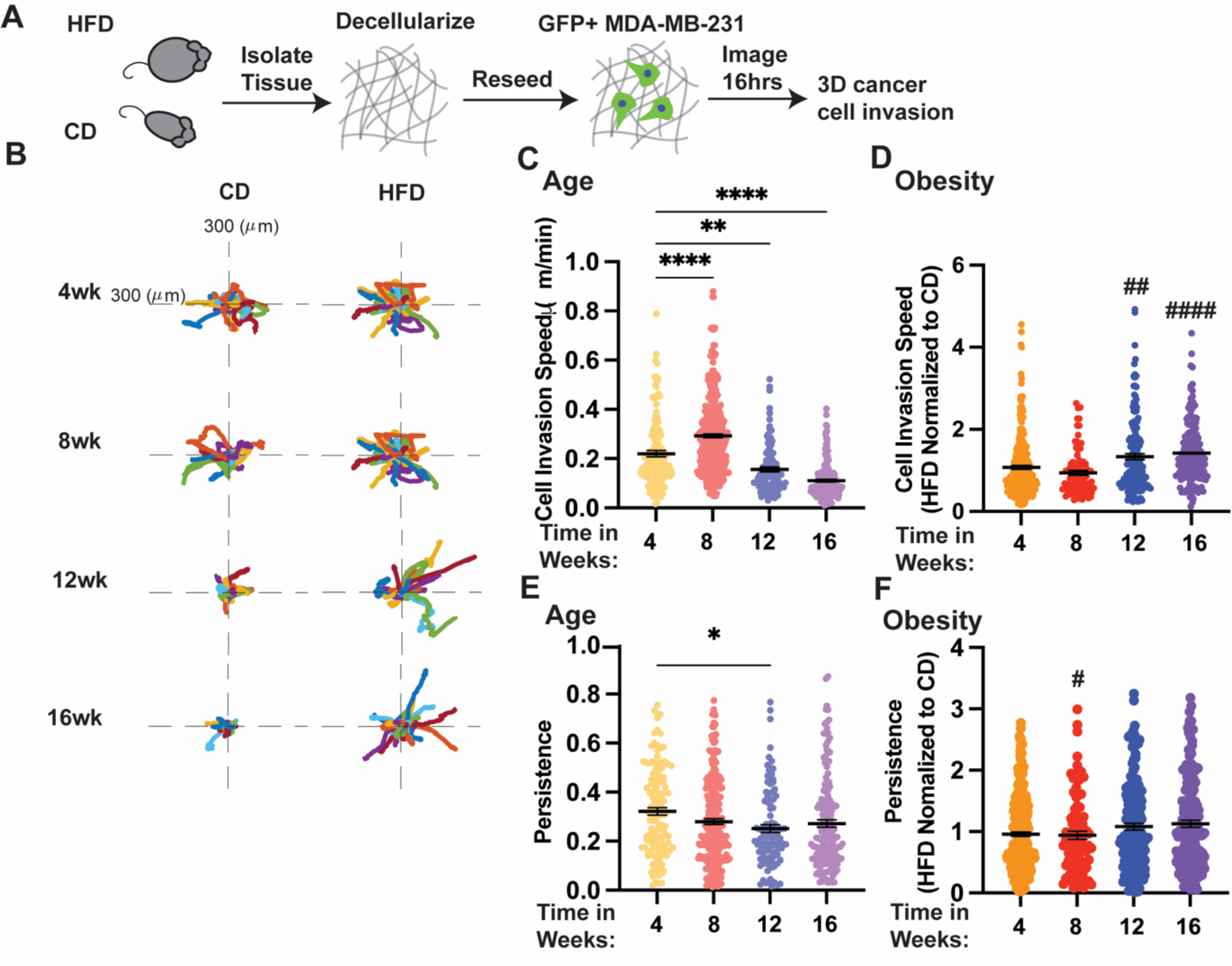
Decellularized mammary fat pad ECM from younger aged mice and later-stage obesity had the highest invasive potential. **(A)** Schematic depicting experimental procedure mammary fat pads were isolated, decellularized using 0.1% SDS solution, reseeded with GFP+ MDA-MD-231 human TNBC cells, and time-lapse imaged. **(B)** Representative rose plots of cell path invading on decellularized ECM from mice fed CD or HFD for 4, 8, 12, or 16 weeks. **(C)** Cell invasion speed of MDA-MB-231 cells invading on decellularized ECM from mice at different ages after being started on CD. **(D)** Cell invasion speed of MDA-MB-231 cells invading HFD relative to CD decellularized ECM scaffolds. **(E)** Persistence of MDA-MB-231 cells invading decellularized ECM from mice at different ages after being started on CD. **(F)** Persistence of MDA-MB-231 cells invading on HFD relative to CD decellularized ECM scaffolds. Significance was determined by ordinary one-way ANOVA with Tukey multiple comparisons to compare between CD groups (P*<0.05, P**<0.01, P***<0.001, P****<0.0001) only comparisons to 4 weeks are shown. An unpaired t-test was used to compare age-matched CD and HFD groups (P#<0.05, P##<0.01, P###<0.001, P####<0.0001). Data are shown as mean with SEM each data point represents a single cell, cells seeded on decellularized ECM scaffolds from at least n=3 mice.

### Decellularized mammary fat pad ECM from younger mice and later-stage obesity have the highest invasive potential

We then sought to understand how aging and DIO tissue-level ECM changes in the mammary fat pad affect TNBC cell motility, a phenotype important for local tumor growth, invasion, and distant metastasis. To do this, we fed non-tumor bearing mice HFD or CD for 4, 8, 12, and 16 weeks, then isolated and decellularized the mammary fat pads to generate ECM scaffolds using our previously published method [23]. Fluorescently labeled GFP+ MDA-MB-231 cells, a human TNBC cell line, were reseeded onto scaffolds, timelapse imaged, and tracked to quantify cell invasion and persistence (Fig 2A, B). We saw a significant age-related increase in the invasion speed of cells reseeded on decellularized ECM at 8 weeks on CD, and then a significant decrease in cell invasion at 12 and 16 weeks (Fig 2B, C). At 12 and 16 weeks on HFD, cell invasion speed was significantly faster in the ECM from HFD-fed mice than in age-matched control (Fig 2 C, D, S3A). Persistence measures the Euclidean distance between the start and finish of the cell’s path over the total distance traveled and has been associated with increased metastasis. Age did not have a consistent effect on the persistence of cells invading decellularized ECM (Fig 2E, S3B). On HFD, there was a significant decrease in persistence at 8 weeks compared to age-matched CD and other time points (Fig 2F, S3B). These data show that decellularized mammary fat pad ECM from younger-aged mice (4 and 8 weeks on CD) and later-stage obese mice (12 and 16 weeks on HFD) had the highest invasive potential.

### Age and obesity induce compositional changes in the mammary fat pad ECM

To identify protein level ECM changes in the mammary fat pad that could be promoting TNBC cell invasion in the context of age and obesity, we performed proteomics on mammary fat pads from female C57BL/6 mice at each time point 4, 8, and 12 weeks on CD or HFD. Principal component analysis on the resulting matrisome composition data revealed clustering of ECM composition at all ages on CD while, distinct differences in the ECM on HFD resulted in separate clustering of each group based on time point (Fig 3A). In the core matrisome of mammary fat pads, we identified 15 collagen chains, 42 glycoproteins, 7 proteoglycans, 39 ECM-regulator proteins, and 20 ECM-affiliated proteins differentially regulated depending on age and diet (Fig S4,5). Comparing mammary fat pad ECM composition on CD at each age from 4 weeks after initiation of diet, 46% of collagens were upregulated at 8 weeks and 93% at 12 weeks on CD (Fig 3B). In comparing mammary fat pad ECM composition on HFD to age-matched CD, 26% of collagens were upregulated after 4 weeks on HFD, 80% after 8 weeks, and 46% after 12 weeks (Fig 3C).

**Figure 3.**
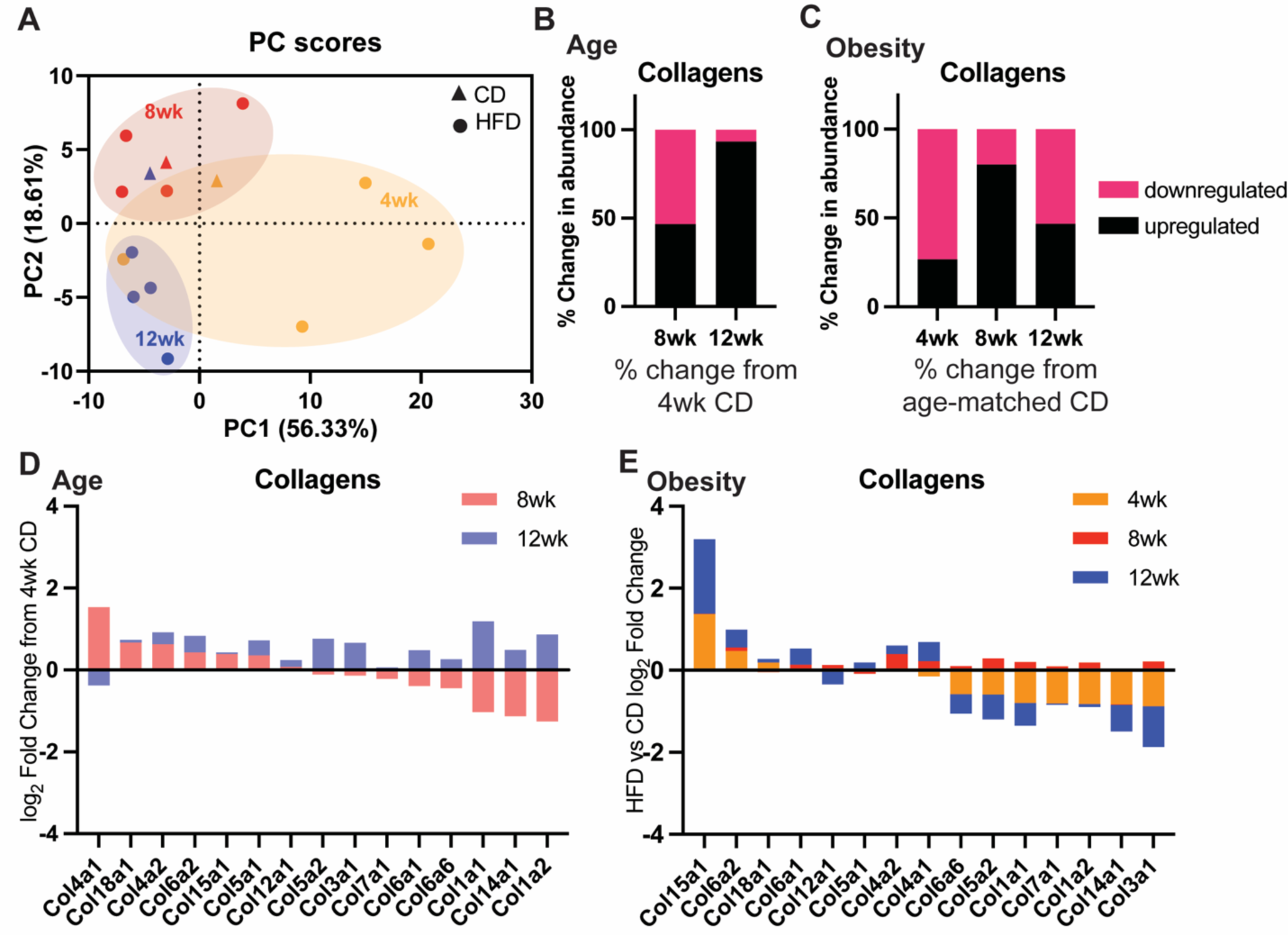
Proteomics analysis identified distinct differences in mammary fat pad ECM at 4, 8, and 12 weeks on HFD. **(A)** Principal component analysis of ECM composition of mammary fat pads at 4, 8, and 12 weeks on CD and HFD. Percent change in abundance or upregulated and downregulated collagens with (**B)** age compared to 4 weeks on CD and **(C)** HFD compared to age-matched CD. **(D)** log_2_ Fold Change in ECM protein abundance in mammary fat pads with age. **(E)** log_2_ Fold Change in mammary fat pads protein abundance at 4, 8, and 12 weeks on HFD relative to age-matched CD.

Collagen IV and VI are both known to be drivers of TNBC invasion [23], [39], [40] and are comprised of three chains that form a triple-helix structure. Collagen IV is comprised of distinct chains ⍺1(IV)-⍺6(IV) that assemble into 3 distinct heterotrimers, ⍺1⍺1⍺2, ⍺3⍺4⍺5, ⍺5⍺5⍺6 [37]. Collagen VI is comprised of chains ⍺1(VI), ⍺2(VI), and a third chain ⍺3(VI)-⍺6(VI) that share a similar structure to ⍺3(VI) to form Collagen VI [38]. After 8 weeks on CD, we see that Col4a1, Col4a2, and Col6a2 were upregulated and Col6a1 was downregulated and at 12 weeks on CD, Col4a2, Col6a2, and Col6a1 were upregulated and Col4a1 was downregulated (Fig 3D). HFD increased the abundance of both Collagen IV and Collagen VI ⍺1 and ⍺2 chains as obesity progresses (Fig 3E).

To quantify full-length Collagen IV and VI in the mammary fat pads, we performed immunostaining on the mammary fat pads. We identified age-dependent enrichment of Collagen IV in the mammary fat pads of CD fed mice, with Collagen IV peaking at 8 weeks and then decreasing after that (Fig 4A, B). We also confirm that there is a significant positive correlation between Collagen IV abundance in the mammary fat pad and total body weight in the context of obesity (Fig 4C). Collagen VI is enriched in the mammary fat pads of mice starting at 12 weeks, and there is a significant positive correlation between Collagen VI abundance in the mammary fat pad and total body weight (Fig 4D-F). These data along with the proteomics data indicate that Collagen IV is initially enriched with age then lowers in abundance and ultimately, plateaus. Conversely, Collagen VI continues to be enriched with age. These data also show that HFD and increasing body weight progressively enriched Collagen IV and VI abundance in the mammary fat pad. Thus, we characterize age-driven and obesity-driven changes in the ECM of the mammary fat pad and identify Collagen IV and VI as proteins whose levels are abundant in conditions where decellularized ECM increased breast cancer cell invasion.

**Figure 4.**
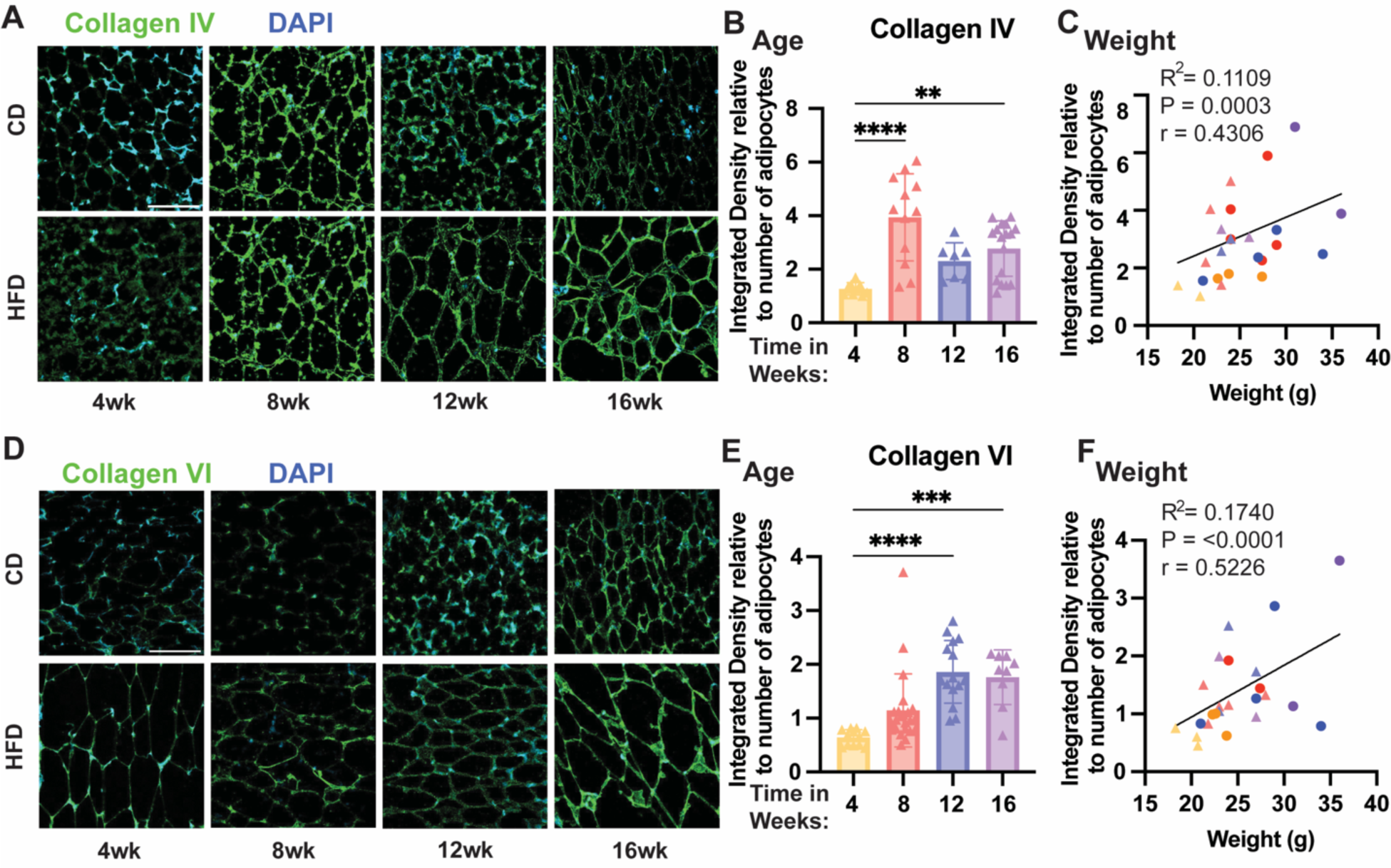
Collagen IV and Collagen VI abundance in mammary fat pads on HFD and CD. **(A)** Representative images of fat pads stained for Collagen IV (green) and DAPI (blue) fed HFD or CD for 4, 8, 12, and 16 weeks. **(B)** Abundance of Collagen IV (integrated density) relative to the number of adipocytes in mice fed CD for 4, 8, 12, and 16 weeks. **(C)** Correlation between abundance of Collagen IV (integrated density) relative to the number of adipocytes and body weight of mouse. (**D)** Representative images of mammary fat pads stained for Collagen VI (green) and DAPI (blue) fed HFD or CD for 4, 8, 12, and 16wks. **(E)** Abundance of Collagen VI (integrated density) relative to the number of adipocytes in mice fed CD over 4, 8, 12, and 16 weeks. **(F)** Correlation between the abundance of collagen VI (integrated density) relative to the number of adipocytes in FOV and mouse body weight. Significance was determined by ordinary one-way ANOVA with Tukey multiple comparisons to compare between CD groups or HFD relative to CD groups (P*<0.05, P**<0.01, P***<0.001, P****<0.0001). Data are shown as mean with SD n = 3 animals/group.

### Cells of the stromal vascular fraction (SVF), not mature adipocytes are responsible for differential ECM deposition in obese adipose tissue

Next, we sought to determine which cell types in the adipose tissue microenvironment are expressing and secreting Collagen IV and Collagen VI. Collagen IV has been shown to be expressed by adipocytes, endothelial cells, fibroblasts, and macrophages in adipose tissue [41]–[44]. We used a published single-cell RNA seq data set to investigate cell expression of COL4A-6 and COL6A1-6 from white adipose tissue in humans categorized as lean (BMI 20-30), overweight (BMI 30-40), and obese (BMI 40-50), as well as mice categorized as lean (CD) and obese (HFD) [48]. COL4 chains are predominately expressed by adipocytes with low expression levels in ASPCs, pericytes, endothelial, lymphatic endothelial cells (LECs), smooth muscle cells, and endometrium (Fig 5A). Interestingly, adipocytes in adipose tissue are primarily implicated in expressing Collagen VI [38], [45]–[47]. In humans, the COL6A4 chain is not functional [45]. COL6 chains in human, white, adipose tissue are predominantly expressed by ASPCs and with low mRNA expression by adipocytes and cells of the vascular and ductal structures, mesothelium, pericytes, smooth muscle cells, and endometrium (Fig 5B). These same cell types expressed Col4a1-6 and Col6a1-6 in mice (Fig S6A).

**Figure 5.**
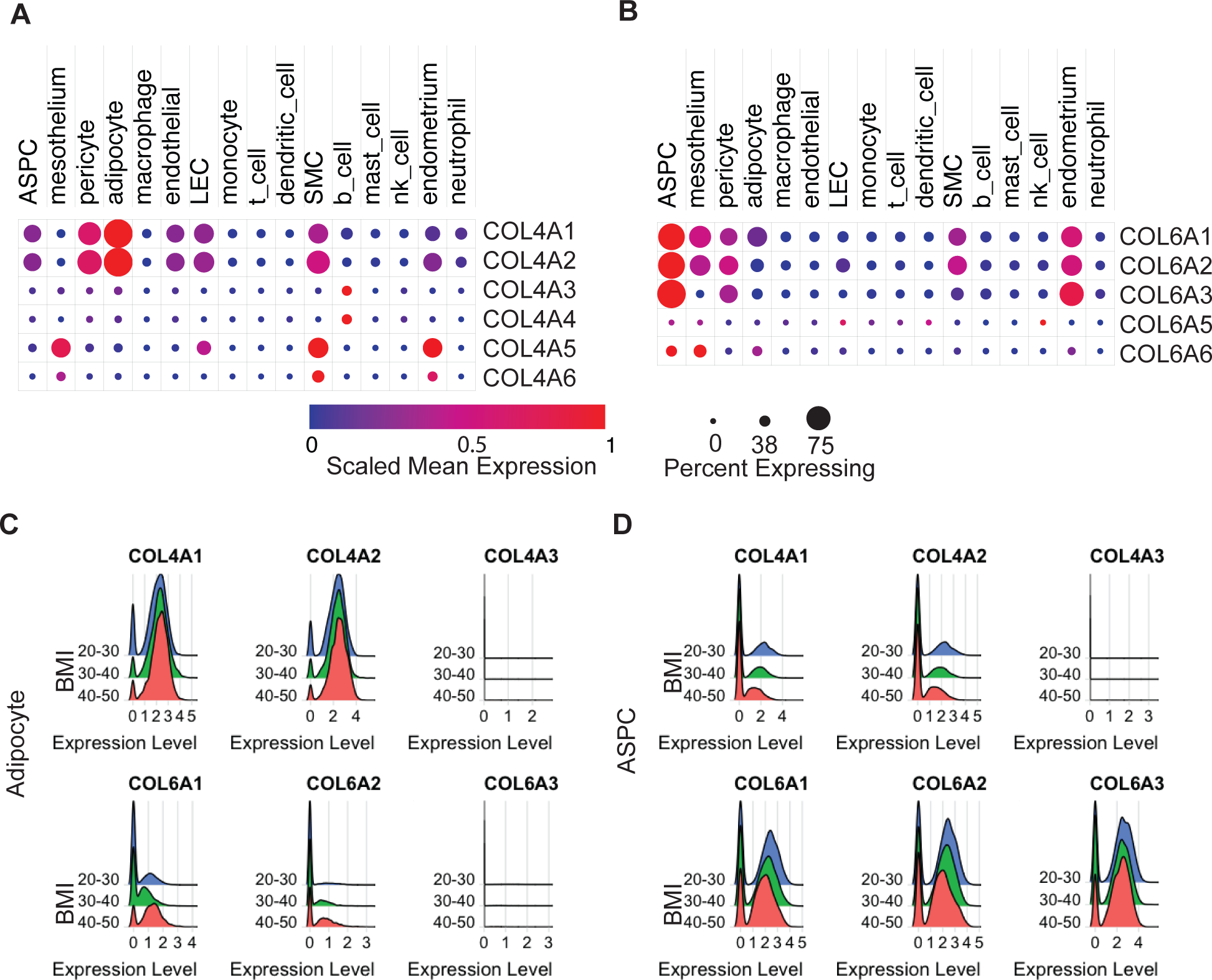
Human RNA-Seq data from human white adipose tissue. Scaled mean expression (blue low red high) and percentage of cells (size of the dot) of human white adipose tissue expressing (**A)** COL4A1-6 and (**B)** COL6A1-6 (excluding COL6A4 not expressed in humans). (**C)** COL4A1-3 and COL6A1-3 expression by female human subcutaneous adipocytes. (**D)** COL4A1-3 and COL6A1-3 expression by female human subcutaneous ASPC. Data from the Broad Institute [48].

Since local TNBC invasion occurs when tumor cells begin to invade the local breast stroma, we looked at Collagen IV and VI expression in the RNA seq data by only subcutaneous adipocytes and ASPCs isolated from females of varying BMI. Mature adipocytes are large round lipid-filled cells in adipose tissue and ASPCs are a part of the SVF that are not immune or vascular cells and differentiate into mature adipocytes [49]. Since there was little to no expression of genes that code Collagen IV or VI ⍺4-⍺6 chains in either human or mouse adipose tissue from the RNA-seq data, we chose to focus only on chains ⍺1-⍺3. In human adipocytes, there was no difference in COL4A1-2 expression in lower versus higher BMIs, and there was minimal COL4A3 expression (Fig 5C). In mouse adipocytes, there was an increase in Col4a1 and Col4a2 in HFD (Fig S6C). In human ASPCs, there was no COL4A3 expression and low expression of COL4A1 and COL4A2 in all BMI groups (Fig 5D). In mouse ASPCs, Col4a1 and Col4a2 were poorly expressed in CD and HFD groups, and Col4a3 was minimally expressed, (Fig S6D). These data indicate that Collagen IV is primarily expressed by adipocytes in adipose tissue in both humans and mice. BMI did not influence expression levels in human adipocytes, but expression levels increased by adipocytes in HFD mice compared to CD mice.

In human adipocytes, we saw an increase in COL6A1-2 expression but minimal COL6A3 expression in the high BMI group (BMI 40-50) (Fig 5C). In mouse adipocytes, we saw no Col6a3 expression and low Col6a1-2 expression in both CD and HFD groups, with slightly higher expression in the HFD group (Fig S6C). In human ASPCs, COL6A1-3 chains were highly expressed independent of BMI in ASPCs (Fig 5D). In mouse ASPCs, Col6a1-3 chains were expressed at higher levels by the HFD group (Fig S6D). These data indicate that in adipose tissue Collagen VI is primarily expressed by ASPCs, and not adipocytes. BMI in humans and diet in mice did not influence expression levels in human ASPCs, but obese BMI in humans and HFD mice correlated with increased expression by adipocytes.

Next, we examined the protein-level secretion of Collagen IV and VI by adipocytes and stromal vascular cells. We isolated subcutaneous mammary fat pads from CD mice, separated the floating adipocyte fraction and a pelleted the SVF containing ASPCs, and quantified Collagen IV and VI protein production *in vitro*. We found that cells of the SVF secreted Collagen IV extracellularly. While Collagen VI could be detected, it was primarily intracellular (Fig 6A). To culture and image the adipocytes and secreted ECM, we employed two cell culture techniques: a ceiling culture technique where a glass cover slip was placed on top of the floating adipocytes, and a 3D gel where adipocytes are encapsulated in Collagen I, the most abundant ECM protein breast tissue. Interestingly, in our ceiling culture, we found extracellular Collagen IV, but minimal Collagen VI (Fig 6B). When adipocytes were encapsulated in Collagen I, we saw minimal Collagen IV and VI in the cultures (Fig 6C). These data demonstrate that stromal vascular cells, specifically ASPCs, and not adipocytes are responsible for secreting Collagen IV and VI ECM in subcutaneous adipose tissue.

**Figure 6.**
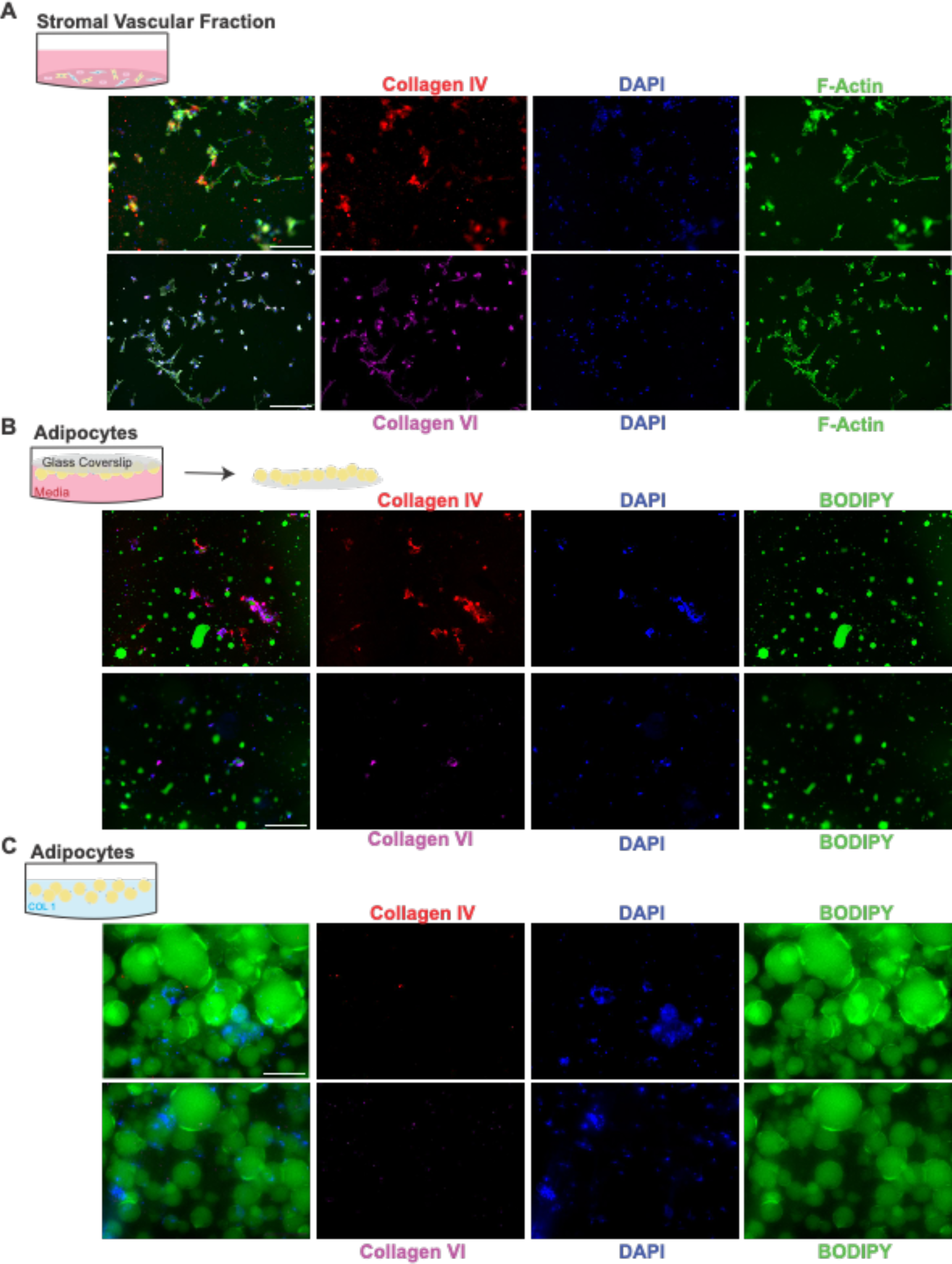
Collagen IV and VI secretion by adipose SVF cells and adipocytes. **(A)** Collagen IV (red) and Collagen VI (magenta), F-actin (green), and DAPI (blue) staining of isolated mouse SVF fraction. (**B)** Isolated mouse adipocyte ceiling cultures and **C)** isolated mouse adipocytes encapsulated in Collagen I stained for Collagen IV (red) and Collagen VI (magenta), BODIPY (green), and DAPI (blue). Scale bar = 200µm.

### HFD reverses age-related reduction in invasive potential of whole decellularized lung and liver ECM

Obesity is a systemic disease that has also been shown to affect the lungs and livers, common sites of TNBC metastasis, through inflammation and lipid accumulation, respectively [18], [25]. We found a positive correlation between weight gain and lung metastasis (Fig 1H). While in the liver, we saw an increase in the number of metastases on HFD compared to CD only at 4 weeks. We next sought to investigate whether obesity-dependent changes in the whole ECM from lungs and livers could be driving TNBC cell invasion, a process important for metastatic outgrowth. We isolated and decellularized lungs and liver from non-tumor bearing, 10-week-old, female mice fed HFD or CD for 4, 8, 12, and 16 weeks, reseeded them with fluorescently labeled human MDA-MB-231 cells, and quantified cell invasion speed and persistence. Older age (16 weeks after initiation of CD) decreased cell invasion speed on decellularized lung ECM, while HFD reversed this age-related effect by significantly increasing invasion speed compared to age-matched CD at 16 weeks on HFD (Fig 7 A-C, S7A). There was no age-related difference in the persistence of tumor cells invading decellularized lung ECM, but HFD increased the persistence at 16 weeks relative to CD (Fig 7D, E S7B). While age hindered the metastatic potential of lung ECM, HFD reversed these effects and promoted cell invasion.

**Figure 7.**
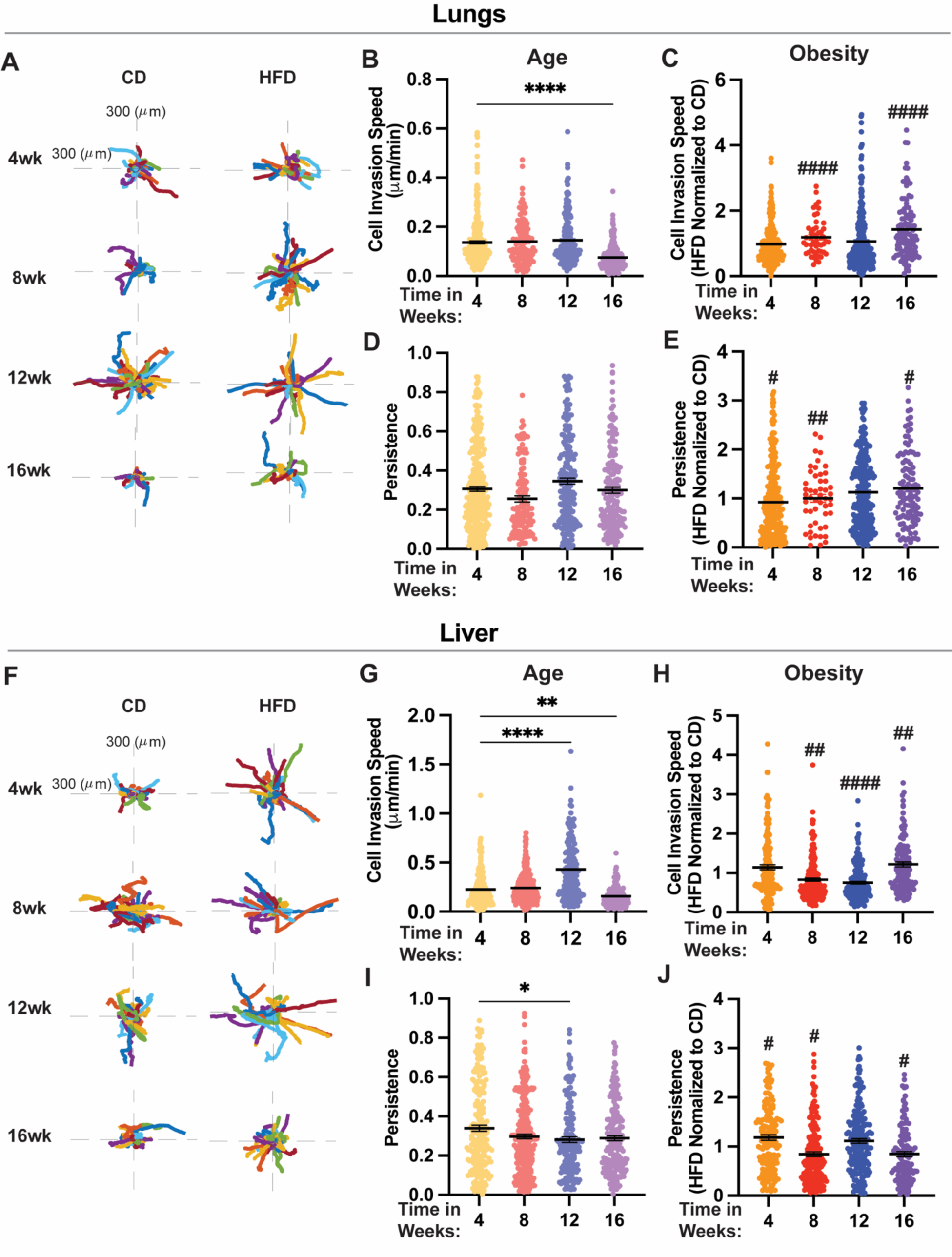
Tumor cell Invasion on decellularized lung ECM. **(A)** Representative rose plots of cell path invading on decellularized lung ECM fed CD or HFD for 4, 8, 12, or 16 weeks, **(B)** cell invasion speed, and **(C)** persistence of MDA-MB-231 cells invading on decellularized lung ECM from CD-fed mice at different ages. **(D)** Cell invasion speed and **(E)** persistence of MDA-MB-231 cells invading on HFD relative to CD decellularized lung ECM scaffolds. **(F)** Representative rose plots of cell path invading on decellularized liver ECM fed CD or HFD for 4, 8, 12, or 16 weeks. **(G)** Cell invasion speed and **(H)** persistence of MDA-MB-231 cells invading on decellularized liver ECM from CD-fed mice at different ages. **(I)** Cell invasion speed and **(J)** persistence of MDA-MB-231 cells invading on HFD relative to CD decellularized liver ECM scaffolds. Significance was determined by ordinary one-way ANOVA with Tukey multiple comparisons to compare between CD groups or HFD relative to CD groups (P*<0.05, P**<0.01, P***<0.001, P****<0.0001). An unpaired t-test was used to compare age-matched CD and HFD groups (P#<0.05, P##<0.01, P###<0.001, P####<0.0001). Data are shown as mean with SD n = 8-10 animals/group. Each data point represents a single cell, cells seeded on decellularized ECM scaffolds from at least n=3 mice.

In the liver, the age of decellularized liver ECM increased the invasion speed of cells on ECM 12 weeks after starting CD then decreased the invasion speed at 16 weeks (Fig 7F, G). HFD decreased invasion speed at 8 and 12 weeks relative to CD and then increased invasion speed at 16 weeks (Fig 7H S8A). Similar to the lungs, HFD for 16 weeks reversed the age-related hindrance on cell invasion and promoted cell invasion. Age decreased the persistence of cells invading on liver ECM at 12 weeks on CD (Fig 7I). HFD reversed this decline in persistence seen at 12 weeks in the CD group, but HFD compared to age-matched CD only significantly increased the persistence of cells on decellularized liver ECM at 4 weeks (Fig 7J, S8B). These data indicate that later stages of HFD reverse the age-related decrease in the metastatic potential of whole ECM. Since little is known about liver metastasis in the context of obesity and breast cancer, and there is confounding evidence for the role of steatosis in promoting liver metastasis, we chose to focus on investigating compositional changes in the liver ECM.

### Enrichment of ECM drivers of invasion in the liver at a younger age and early DIO

Obesity affects the liver by causing lipid accumulation in hepatocytes, a process known as steatosis. We isolated livers from healthy, 10-week-old female mice fed CD or HFD for 4, 8, 12, and 16 weeks, weighed, fixed, and stained them with H&E. We found no difference in liver weight with age or HFD (Fig S9A). Starting at 12 weeks on HFD H&E staining shows evidence of macrosteatosis (Fig S9B). We conducted proteomic analysis on liver ECM from mice fed HFD or CD for 4, 8, and 12 weeks to investigate age and early obesity-dependent changes in ECM. Principal component analysis on the resulting matrisome composition data identified distinct differences in the ECM composition of the liver at all ages on CD and distinct differences at 4, 8, and 12 weeks on HFD, resulting in the clustering of each group (Fig 8A). We identified 14 collagens in the core matrisome of the livers. In comparing liver ECM composition at each age from 4 weeks after initiation of CD, 36% of collagens were upregulated at 8 weeks and 29% at 12 weeks on CD (Fig 8B). Comparing mammary liver ECM composition on HFD to age-matched CD, 64% of collagens were upregulated after 4 weeks, 50% after 8 weeks, and 35% after 12 weeks (Fig 8C). In the core matrisome, we identified 17 glycoproteins and 5 proteoglycans, along with 32 ECM-regulator proteins, and 17 ECM-affiliated proteins differentially regulated with age and HFD (Fig 8 E-G, S10, 11).

**Figure 8.**
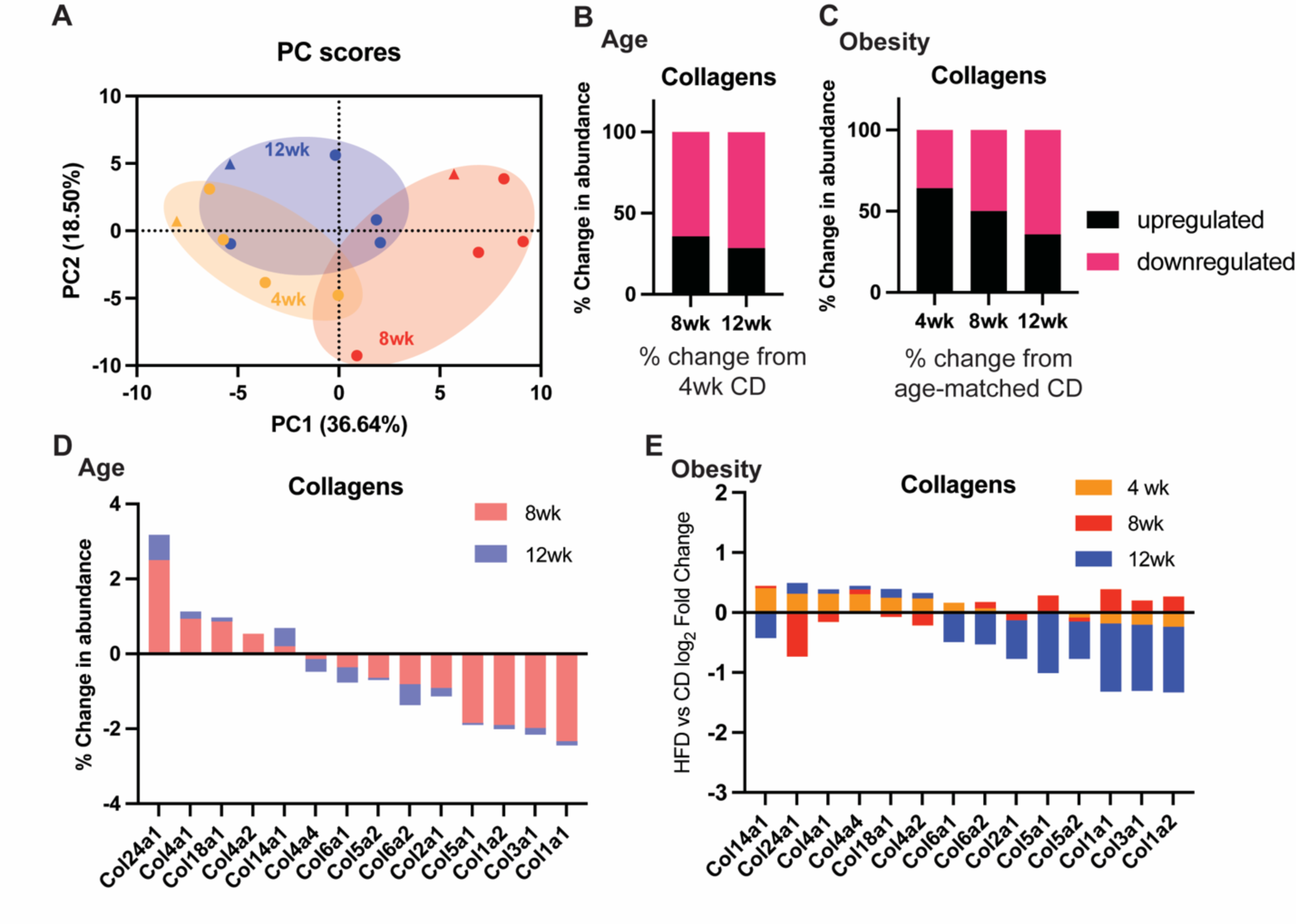
Proteomics analysis identified distinct differences in liver ECM with age and at 4, 8, and 12 weeks on HFD. **(A)** Principal component analysis of ECM composition of mammary fat pads at 4, 8, and 12 weeks on CD and HFD. Percent change in abundance or upregulated and downregulated collagens with (**B)** age compared to 4 weeks on CD and **(C)** HFD compared to age-matched CD. **(D)** log_2_ Fold Change in ECM protein abundance in mammary fat pads with age. **(E)** log_2_ Fold Change in mammary fat pads protein abundance at 4, 8, and 12 weeks on HFD relative to age-matched CD.

Interestingly, we found upregulation of Col4a1-2 and downregulation of Col5a1 and Col6a1-2 with age. Collagen V in the liver has been shown to be an important component of the pre-metastatic niche that promotes metastatic outgrowth [50]. We also see enrichment of collagen chains that are drivers of invasion, Col4, Col5, and Col6, at 4 weeks on HFD, but that are downregulated at later time points on HFD (Fig 8D,E). These data indicate that early stages of obesity are sufficient to induce tissue-level changes in the liver ECM that drive TNBC invasion and metastasis in the liver early on in DIO but not at later stages.

## Discussion

Obesity is a growing public health crisis known to be a risk factor for a range of diseases including breast cancer. In fact, the rate of breast cancer incidence in women has been increasing by about 0.5% per year, which is partly attributed to increases in excess body weight [51]. Furthermore, obesity is a risk factor for chemoresistance and increased metastasis [5], [24]. Breast cancer in younger women is associated with more aggressive disease and lower survival rates, with TNBC being the most common subtype of breast cancer among younger women [52]. Thus, there is a critical need to identify potential age and obesity-dependent drivers of TNBC progression to better diagnose and treat these patient populations. Overall, our combined data show that age had no effect on primary tumor growth, and younger age increased ECM-driven invasion while older age increased metastasis to the lungs and liver, but decreased ECM-driven invasion. On the other hand, obesity increased primary tumor growth and mammary fat pad ECM-driven invasion at 12 weeks, and weight gain correlated to increased lung metastasis while liver metastasis only increased early in DIO (4 weeks). In the mammary fat pad, we found enrichment of Collagen IV and Collagen VI as obesity progressed and identified cells of the SVF and ASPCs (not adipocytes) as the cell types primarily responsible for secreting Collagen IV and VI in the breast adipose tissue. Together, these data demonstrate that both age and obesity contribute to tumor and metastatic outgrowth and implicate ECM changes in the primary tumor and metastatic microenvironment as a mechanism by which age and obesity impact breast cancer progression.

In humans, obesity is a heterogeneous disease, due to variables such as sex, age, and duration of obesity [53]. Few studies using murine models have explored this heterogeneity in the context of breast cancer, which we tackled by characterizing mammary tumor growth, metastasis, and the ECM in mammary adipose tissue at different time points representing the progression of age and obesity. In our murine DIO model at 4 weeks on HFD there is a significant increase in total body weight, mammary fat pad weight, and adipocyte size, with few effects on tumor growth and ECM-driven invasion, but increased liver metastasis. By 8 weeks on HFD, we can start to see significant changes in the abundance of ECM proteins produced by ASPCs that drive tumor cell invasion in the mammary gland. By 12 weeks on HFD, there was a significant increase in primary tumor growth. With age alone, we also saw a significant increase in body weight starting at 12 weeks on CD. Notably, this study highlights the importance of consistency when using DIO murine models in studying complex diseases. As small as a few-week difference in age or duration of diet has a significant impact on adipose tissue physiology and TNBC progression that could influence results from testing drugs and treatments in diverse models of age and obesity.

Patients with TNBC whose tumors have invaded the breast adipose tissue microenvironment have a significantly poorer prognosis than those without invasion which is further exacerbated by obesity [54]. Adipose tissue is comprised of adipocytes encapsulated in ECM that are remodeled to accommodate adipocyte hyperplasia and hypertrophy during weight gain. Collagen IV is well characterized to be secreted by cells found in the adipose SVF, fibroblasts, macrophages, and endothelial cells [42], [55]. Our study raises an interesting question about what cell types are involved in the expression and secretion of Collagen VI in adipose tissue. The literature primarily implicates adipocytes as the primary cell type in adipose tissue secreting Collagen VI [38], [40]. However, using an RNA-Seq dataset and primary cell cultures we show that isolated mature adipocytes express and secrete low levels of Collagen VI, with ASPCs having much higher expression than adipocytes in both mice and humans. This discrepancy between our data and the literature could be due to the lack of expression of the Col6a3 chain by adipocytes preventing the formation of assembled Collagen VI monomers. Additionally, the majority of studies investigating adipocyte-secreted Collagen VI use differentiated mouse 3t3L-1 fibroblasts and human bone marrow-derived stem cells [40], [45]–[47]. One study found that in 3t3L-1s Collagen IV and VI secretion peaked during differentiation [55]. Similar to our findings, one study by McCulloch et al. using primary isolated adipose cells found that there was a 5-fold higher expression of Col6a3 in SVF cells compared to adipocytes. Thus, enrichment of Collagen IV and VI in obese adipose tissue may be due to adipocyte hyperplasia and the differentiation of progenitor cells to adipocytes as adipose tissue expands, not due to the dysregulation of mature adipocytes. ECM secreted by adipose stromal cells isolated from mammary fat pads of obese versus lean mice has been shown to produce a greater abundance of fibronectin and collagen cross-linking, leading to increased breast cancer cell invasion via mechanosignaling [21]. Thus, adipocytes might play a less important role in ECM remodeling during obesity and cancer than previously thought, with adipose stromal vascular cells-comprised of fibroblast-derived preadipocytes, endothelial cells, and immune cells-playing a more significant role.

The microenvironment of the metastatic site has been shown to be an important contributor to metastatic outgrowth and survival [57]. Once tumor cells leave the vasculature, they enter the new microenvironment at the distant metastatic site where they need to anchor, proliferate, and invade to survive all processes triggered through ECM engagement [57]. Quail et al., investigated the effect of obesity on lung inflammation by quantifying changes in immune cell proportions in the lungs. However, they did not investigate other effects of inflammation and immune cell recruitment such as ECM remodeling. We found that in our DIO model at 16 weeks whole decellularized lung ECM significantly increased tumor cell invasion while an increase in lung metastasis was found at 8 weeks on HFD, indicating that the increase in lung metastasis on HFD relative to CD at earlier time points of obesity may be due to other changes in the lung microenvironment or local mammary adipose tissue that are not related to the lung ECM. Further study could be done to characterize immune cell proportions at earlier stages of obesity since Quail et al. used mice on an HFD for 15 weeks and the link between obesity-driven lung inflammation and ECM changes in the lung may contribute to increased metastasis. Interestingly, there are discrepancies in the literature on the association of liver steatosis with breast cancer metastasis to the liver in patients [33], [34]. We found that at early stages of obesity before steatosis was present there was a significant increase in liver metastasis on HFD but at intermediate and later time points when steatosis was present there was no difference in liver metastasis on HFD compared to CD. In our DIO model, we found that there was no difference in the weight of livers at any stage of obesity, but we saw progressive lipid accumulation forming in the liver starting at 12 weeks. One study found that more advanced stages of obesity (34 weeks on HFD) were required to see an increase in liver weight. In that same study, progressive lipid accumulation in the liver at earlier time points (12 weeks on HFD) was observed [32]. In characterizing the liver ECM, we found that whole decellularized ECM at intermediate stages of obesity (8 and 12 weeks) hindered the invasive potential of the ECM compared to early stages of obesity (4 weeks). We also saw ECM drivers of invasion, Collagen IV, Collagen V, and Collagen VI in the liver decreased in abundance at intermediate stages of obesity. These findings suggest that HFD may induce changes in the liver ECM that inhibit the formation of liver metastasis.

There have been limited mechanistic studies exploring the association between age and TNBC. Interestingly, TNBC has been shown to be more aggressive in young mice than old mice due to differences in inflammation and immune cell activation [58], [59]. However, these studies investigated young mice (6-8 weeks in age) and old mice (>10 months in age) and only quantified primary tumor growth and not metastasis. While we did not see an age-related difference in primary tumor growth, we did see an increase in lung metastasis in mice injected with tumors 12 and 16 weeks after starting on CD (18 and 22-week-old mice) compared to earlier time points (10 and 14-week-old mice). In the liver, we saw an effect where metastasis was increased in mice injected after 4 weeks on diet (10-week-old mice), decreased at 8 and 12 weeks (14 and 18-week-old mice) then increased again at 16 weeks (22-week-old mice). Using our decellularization technique, we identified age-related changes in whole decellularized mammary fat pad, lung, and liver ECM. Our results showed that older age decreased the invasive potential of mammary fat pad ECM and lung ECM. Older age increased the metastatic potential of liver ECM at 12 weeks then followed by a significant decrease at 16 weeks. Proteomics and staining of the mammary fat pad identified an age-related increase in the abundance of Collagen IV and VI. Proteomics of the liver ECM identified distinct age-related compositional differences and a decrease in ECM drivers of tumor cell invasion with age. These findings demonstrate even incremental aging-4 weeks apart-can have a significant impact on metastasis and induce significant changes in the ECM microenvironment at the metastatic sites.

Overall, these findings demonstrate that age and obesity cause significant changes in the microenvironment of not only the primary tumor but the metastatic sites in the lungs and liver. Specifically, age and obesity can both induce changes in the abundance of ECM proteins that contribute to breast tumor cell invasion. In the mammary fat pad, an increase in the abundance of these ECM drivers as obesity progresses promotes tumor cell invasion, tumor growth, and metastasis, while in the liver a decrease in the abundance of these ECM drivers prevents metastasis. Understanding how obesity progresses and how it influences breast cancer progression is important for understanding these complex diseases and identifying potential biomarkers and therapeutic targets for the growing overweight population that is at high risk for developing a range of cancers.

## Methods

### Antibodies, reagents, etc

Antibodies used anti–collagen VI (ab199720; Abcam, Cambridge, MA), and anti-collagen IV (ab6586; Abcam, Cambridge, MA). ECM substrates used were collagen VI (ab7538; Abcam, Cambridge, MA), collagen I (CB-40236; Fisher Scientific, Hampton, NH), Collagen IV protein (ab7536; Abcam, Cambridge, MA)

### Cell Culture

MDA-MB-231 and E0771 cells were obtained from ATCC (Manassas, VA). MDA-MB-231 cells were cultured in Dulbecco’s Modified Eagle’s Medium (DMEM) with 10% serum and Pen-Strep Glutamine. E0771 cells were cultures in Dulbecco’s Modified Eagle’s Medium (DMEM) with 10% serum and Pen-Strep Glutamine and 20 mM HEPES. Cells were checked every 2 months for the presence of mycoplasma by a polymerase chain reaction (PCR) based method using a Universal Mycoplasma Detection

### Animal studies

Female C57Bl/6 mice were purchased from Jackson Laboratories (Bar Harbor, ME) at 8 weeks old. Mice were fed either a high-fat diet with 60kcal% fat (Research Diets, D12492) or control diet with 10kcal% fat (Research Diets, D12450J) for 4, 8, 12, or 16 weeks. Mice were kept on a 12hr light/dark cycle with ad libitum access to food and water. Body weights were measured weekly and at the end of the study and mice were euthanized with carbon dioxide asphyxiation and cervical dislocation. Subcutaneous adipose tissue, livers, and lungs were excised. To model metastatic breast cancer, E0771 murine adenocarcinoma cells (500,000 cells per mouse in phosphate-buffered saline (PBS) and 20% collagen I) were injected into the 4^th^ left mammary fat pad of mice fed either control or high-fat diet for 4, 8, 12, and 16 weeks. Tumor burden was measured weekly with calipers, after 5 weeks or when the maximum tumor burden of 2cm^3^ was reached the animals were euthanized and the tumors, lungs, and livers were excised.

### Decellularized ECM Scaffold Reseeding

Mammary fat, lung, and liver-derived decellularized dECM scaffolds were produced as described previously [22]. Briefly, tissues were dissected from EO771 mice and then were submerged in 0.1% SDS in PBS (w/v) solution with agitation for 3 to 5 days, replacing the solution once daily. Once the tissue had turned completely white, tissues were moved to 0.5% Anti-Anti in PBS solution for 1 day, followed by washing in dH2O for 48 hours to remove residual detergents. Five-millimeter pieces of decellularized scaffold were cut and conditioned in complete cell culture media for 24 hours. Scaffolds were reseeded with 250,000 231-GFP cells and incubated at 37°C for 6 hours. Reseeded tissue was then moved to fresh media, stabilized, and imaged for 16 hours, capturing images every 20 minutes using a Keyence BZ-X710 microscope (Keyence, Elmwood Park, NJ). Cells were then tracked using VW-9000 Video Editing/Analysis Software (Keyence) based on movement of the main cell body, and speed was calculated using a custom MATLAB script vR2018a (MathWorks). Data presented are the result of at least three independent experiments with five fields of view imaged per experiment and 4 to 8 cells tracked per field of view.

### LC-Ms/Ms analysis

Samples were prepared as described previously [22]. Briefly, dECM homogenate was denatured in 8 mol/L urea and 10 mmol/L dithiothreitol, alkylated with 25 mmol/L iodoacetamide, and deglycosylated with peptideN-glycosidase F (P0704S; New England Biolabs). Samples were then digested sequentially, first with endoproteinase LysC (125-05061; Wako Chemicals USA), then trypsin (PR-V5113; Promega). Samples were acidified with 50% trifluoroacetic acid and labeled with TMT10plex (90110; Thermo Fisher Scientific) according to the manufacturer’s instructions. Labeled peptides were fractionated via highpH reverse-phase high-performance liquid chromatography (Thermo Easy nLC1000; Thermo Fisher Scientific) using a precolumn (made in house, 6 cm of 10-mmol/L C18) and a self-pack 5-mmol/L tip analytical column (12 cm of 5-mmol/L C18; New Objective) over a 140-minute gradient before nanoelectrospray using aQExactive HF-X mass spectrometer (Thermo Fisher Scientific). Solvent A was 0.1% formic acid, and solvent B was 80% ACN/0.1% formic acid. The gradient conditions were 2%to 10% B (0 to 3 minutes), 10% to 30% B (3 to 107 minutes), 30% to 40% B (107 to 121 minutes), 40% to 60% B (121 to 126 minutes), 60% to 100% B (126 to 127 minutes), 100% B (127 to 137 minutes), 100% to 0% B (137 to 138 minutes), and 0% B (138 to 140 minutes), and the mass spectrometer were operated in a data dependent mode. The parameters for the full-scan MS were resolution of 60,000 across 350 to 2,000 mass/charge ratio (m/z), automatic gain control 3 × 106, and maximum ion injection time of 50 milliseconds. The full-scan MS was followed by tandem mass spectrometry (MS/MS) for the top 15 precursor ions in each cycle with a normalized collision energy of 34 and dynamic exclusion of 30 seconds.

### Staining and Immunohistochemistry of tissue sections

Mammary fat pads, lungs, and livers dissected from C57BL/6 mice and were fixed in 10% buffered formalin and embedded in paraffin. For H&E staining, standard procedures were followed for H&E, tissue sections were deparaffinized, hydrated, stained with hematoxylin (GHS280; Sigma-Aldrich, St. Louis, MO), and counterstained with eosin (HT110180; Sigma-Aldrich, St. Louis, MO). Stained sections were mounted in toluene (SP15–500,Thermo Fisher Scientific). For Picrosirius red staining, tissue sections were deparaffinized, hydrated, and staining was performed using the Picro Sirius Red Stain Kit (ab150681; Abcam, Cambridge, MA) according to the manufacturer’s instructions, including staining with Picrosirius red solution for 1 hour and washing with 0.5% acetic acid. Sections were mounted with Permount mounting medium (SP15-500, Fisher Scientific, Hampton, NH). For immunofluorescence, tissue sections were deparaffinized followed by antigen retrieval using Citra Plus solution (HK057; Biogenex, Fremont, CA). After blocking in PBS–0.5% Tween 20 and 10% serum, sections were incubated with primary antibodies overnight at 4°C and fluorescently labeled secondary antibodies at room temperature for 2 hours. 4′,6-Diamidino-2-phenylindole (DAPI; D1306; Thermo Fisher Scientific, Waltham, MA) was used to stain cell nuclei, and fluorochromes on secondary antibodies included Alexa Fluor 647 (Jackson ImmunoResearch, West Grove, PA). Sections were mounted in Fluoromount mounting medium (00-4958-02; Thermo Fisher Scientific, Waltham, MA) and imaged using a Keyence BZ-X710 microscope (Keyence, Elmwood Park, NJ) or the Zeiss LSM 900 confocal microscope.

### Adipocyte isolations and culture

Adipocytes and stromal vascular (SVF) cells were isolated from subcutaneous adipose tissue from the mammary fat pad. Isolate adipose tissue was washed with Krebs Ringer bicarbonate buffer (KRBH) (125 mM NaCl, 4.8 mM KCl, 0.5 mM NaH_2_PO_4_, 1.2 mM MgSO_4_, 2.6 mM CaCl_2_, 25 mM Hepes, 2 mM NaHCO_3_, 5.5 mM glucose, 1 % BSA in ultrapure water). The tissue was finely minced using sterile scissors and placed in a solution of KRBH with 1mg/ml collagenase. The tissue was then left to digest at 37°C for 20–30 min with gentle shaking. After digestion, the digested tissue suspension will be filtered with a sterile 200 μm filter mesh on a 50 mL falcon tube. 5–10 mL of medium DMEM/F12 was added to wash the filter. The solution was then centrifuged at 50 × *g* for 5 min and the floating adipocytes were removed. The remaining pelleted stromal vascular fraction was resuspended in DMEM/F12 with 10% FBS and plated on glass bottom dishes for 5 days. The floating adipocytes were washed 3 more times and then plated. Adipocytes were resuspended in α-MEM/F12 plus Insulin plus FBS media. For ceiling cultures, a sterile glass cover slip was placed on resuspended adipocytes and cultured for 5 days. For adipocytes embedded in collagen gels adipocytes were incorporated into a 1mg/mL collagen I (354236; Corning, Corning, NY), 10mM NaOH, 7.5% 10X DMEM, and 50% adipocyte culture media and cultured for 5 days.

### Immunohistochemistry of isolated cells

Then SVF cells and ceiling cultures were fixed using 4% PFA for 10 min adipocytes encapsulated in Collagen I were fixed for 30 minutes. Cells were permeabilized with 0.2% triton-X washed and then blocked with 5% normal donkey serum, and 0.1% tween-20 overnight at 4C. Primary antibodies were then added in a 1% normal donkey serum and 0.1% triton x solution and incubated overnight at 4C. Secondaries including Phalloidin (F-actin) in SVF cells, BODIPY (lipids) in adipocytes, and DAPI (nuclei) were then added in a 1% normal donkey serum and 0.1% triton x solution. Cells were then washed and imaged using a Keyence BZ-X710 microscope (Keyence, Elmwood Park, NJ).

### Human and mouse RNA-seq

For analysis of collagen IV and VI expression in subcutaneous adipose tissue RNA-seq data was explored using the broad institute’s single cell portal to search for gene expression. Adipocytes and ASPCs single cell RNA-seq data was downloaded from the broad institute’s single cell portal and analyzed using the Seurat package in R studio [60]. From this data we selected for cells from only subcutaneous adipose tissue from females [48].

### Statistical analysis

Statistical analysis and visualization were performed using Graph-Pad Prism v8.4.3. To compare two groups, an unpaired Student t-test was used. To compare between more than two groups, a one-way ANOVA was performed using a Tuckey’s multiple testing correction. A P value ≤ 0.05 was considered statistically significant. For proteomics analysis, principal component analysis was performed using Graph-Pad Prism.

## Supplemental Data

**Supplemental Figure 1.**
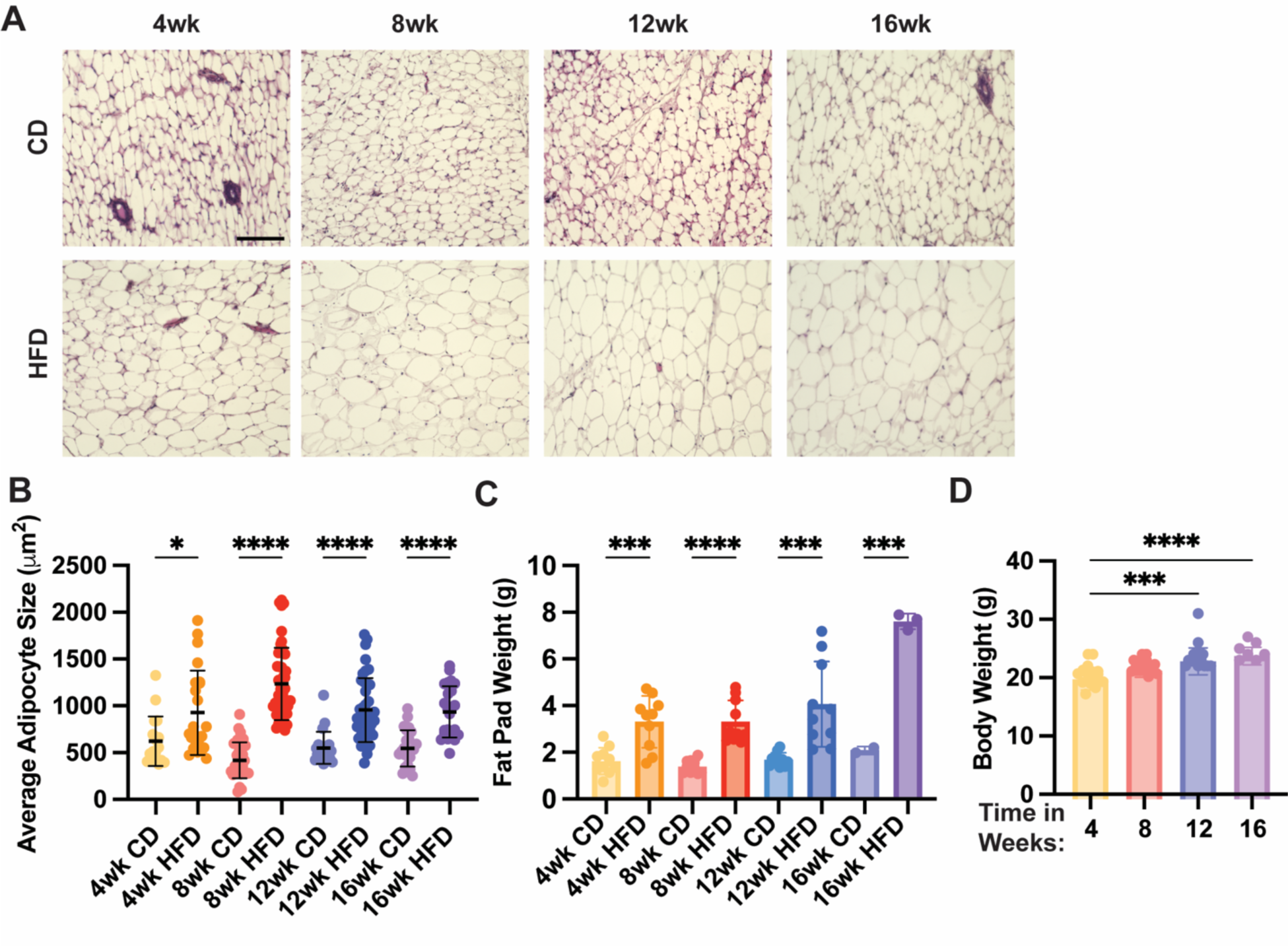
Characterization of mice fed HFD or CD. **(A)** H&E staining of mammary fat pad tissue (Scale bar = 200μm). (**B)** Average size of adipocytes and **(C)** weight of dissected fat pads from mice fed CD or HFD for 4, 8, 12, and 16 weeks. **(D)** Total body weight of mice fed CD for 4, 8, 12, and 16 weeks. (P*<0.05, P**<0.01, P***<0.001, P****<0.0001). Significance was determined using an unpaired t-test to compare age-matched CD and HFD groups and Ordinary one-way ANOVA with Tukey’s multiple comparisons to compare between CD groups. Data are shown as mean with SD n = 3-10 animals/group.

**Supplemental Figure 2.**
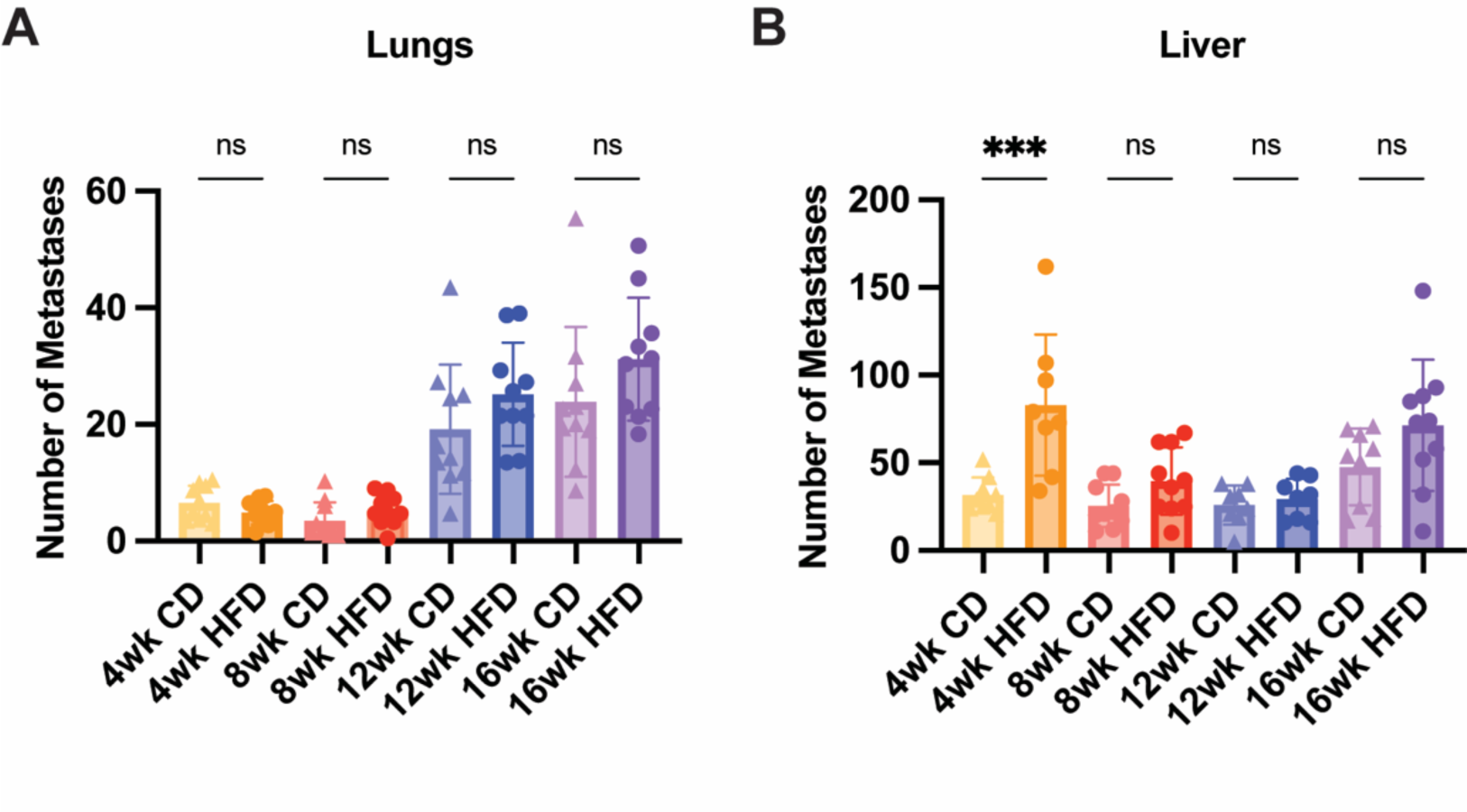
Age-matched CD and HFD lung and liver metastases. **(A)** Age-matched number of lung mets and **(B)** Age-matched number of liver mets at endpoint 5 weeks after tumors were injected in mice fed HFD or CD for 4, 8, 12, and 16 weeks. Significance was determined by an unpaired t-test to compare age-matched CD and HFD groups(P***<0.001). Data are shown as mean with SD n = 8-10 animals/group.

**Supplemental Figure 3.**
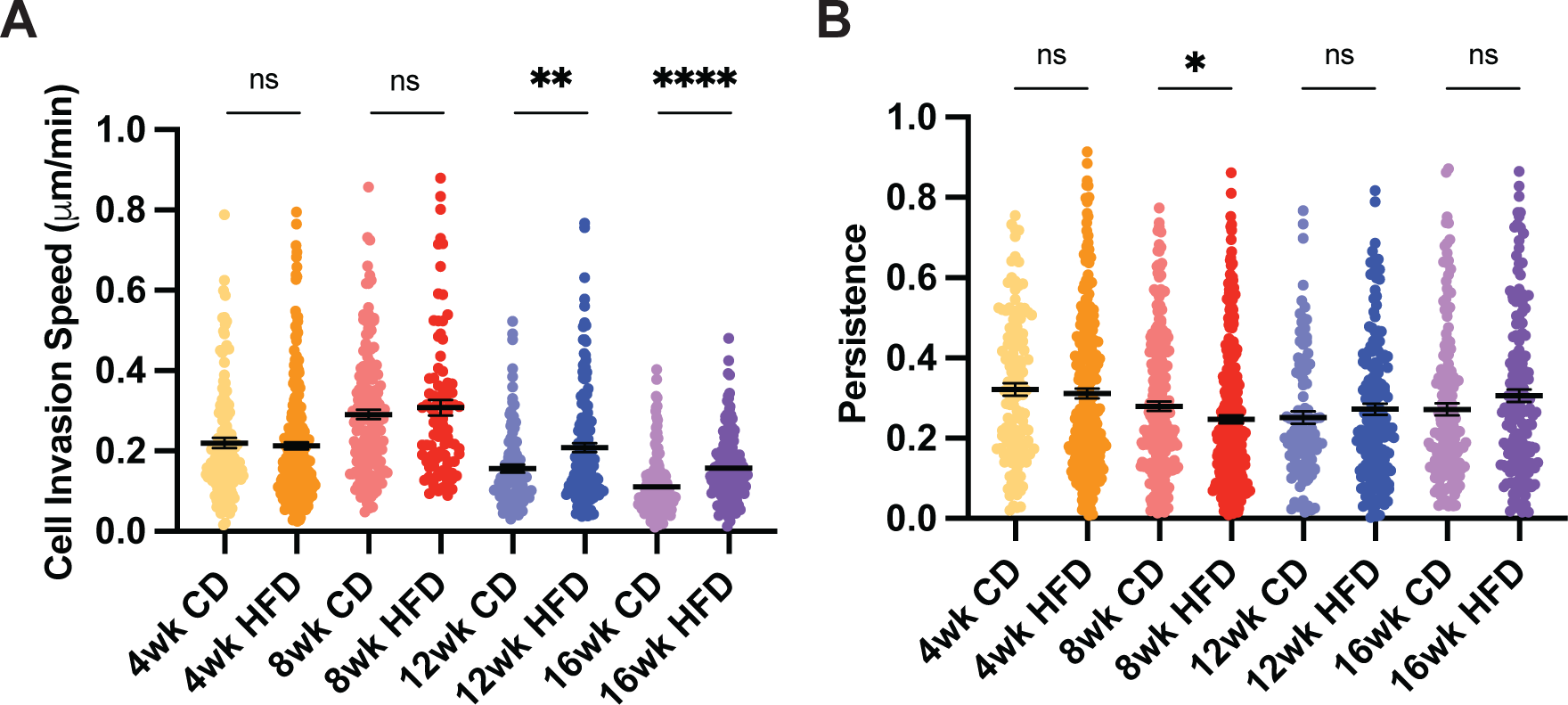
Age-matched CD and HFD cell invasion on mammary fat pad ECM. **(A)** Cell invasion speed and **(B)** persistence of MDA-MB-231 cells invading on decellularized ECM from age-matched CD and HFD-fed mice for 4, 8, 12, and 16 weeks on diet. An unpaired t-test was used to compare age-matched CD and HFD groups (P*<0.05, P**<0.01, P***<0.001, P****<0.0001)

**Supplemental Figure 4.**
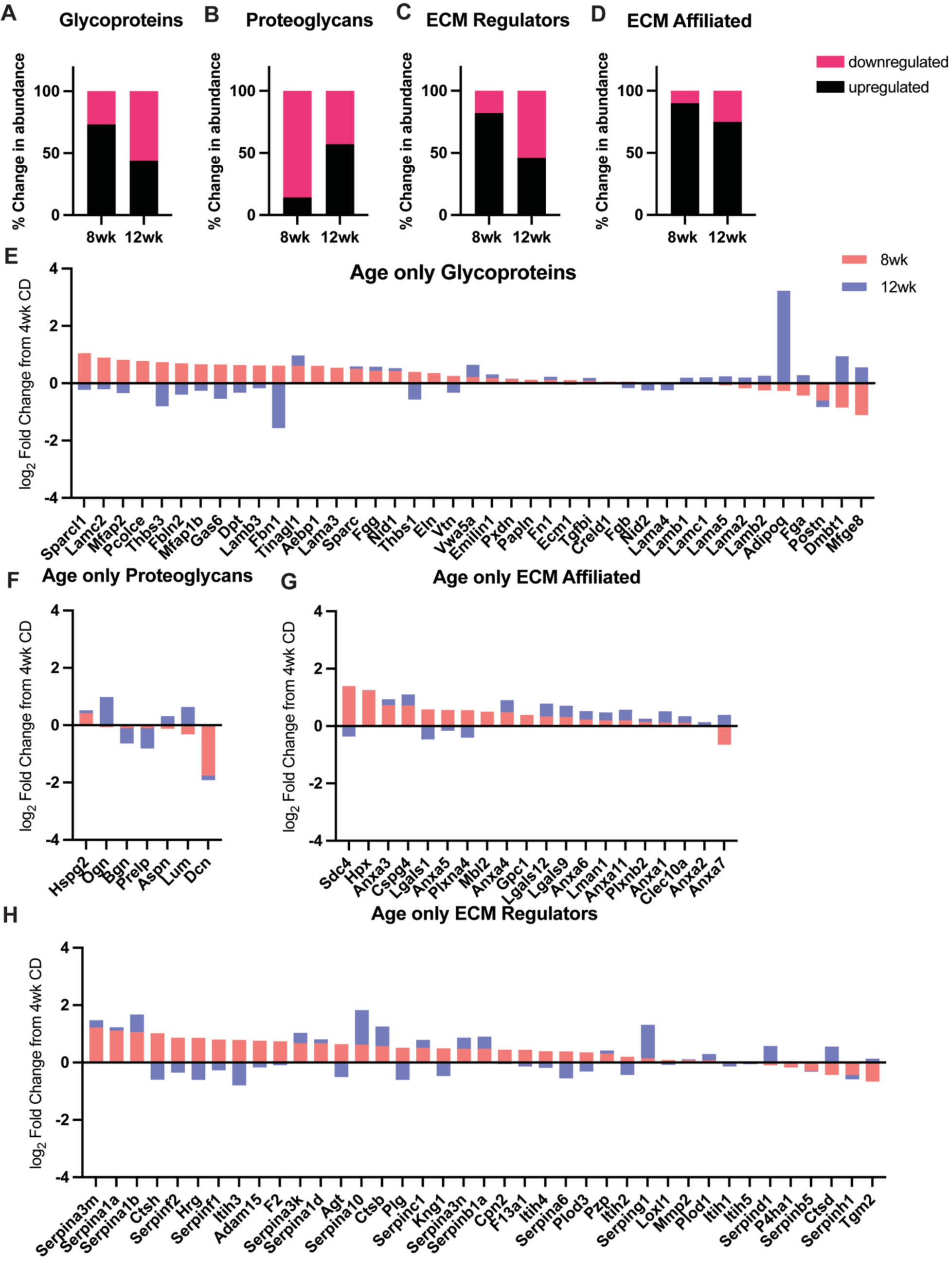
Relative abundance of glycoproteins, proteoglycans, ECM regulators, and ECM-affiliated proteins in the mammary fat pad with age. Percent change in abundance of upregulated and downregulated ECM proteins with age compared to 4 weeks on CD **(A)** glycoproteins, **(B)** proteoglycans, **(C)** ECM regulators, and **(D)** ECM-affiliated. log_2_ Fold Change in ECM protein abundance in mammary fat pads with age compared to 4 weeks on CD **(E)** glycoproteins, **(F)** proteoglycans, **(G)** ECM regulators, and **(H)** ECM-affiliated.

**Supplemental Figure 5.**
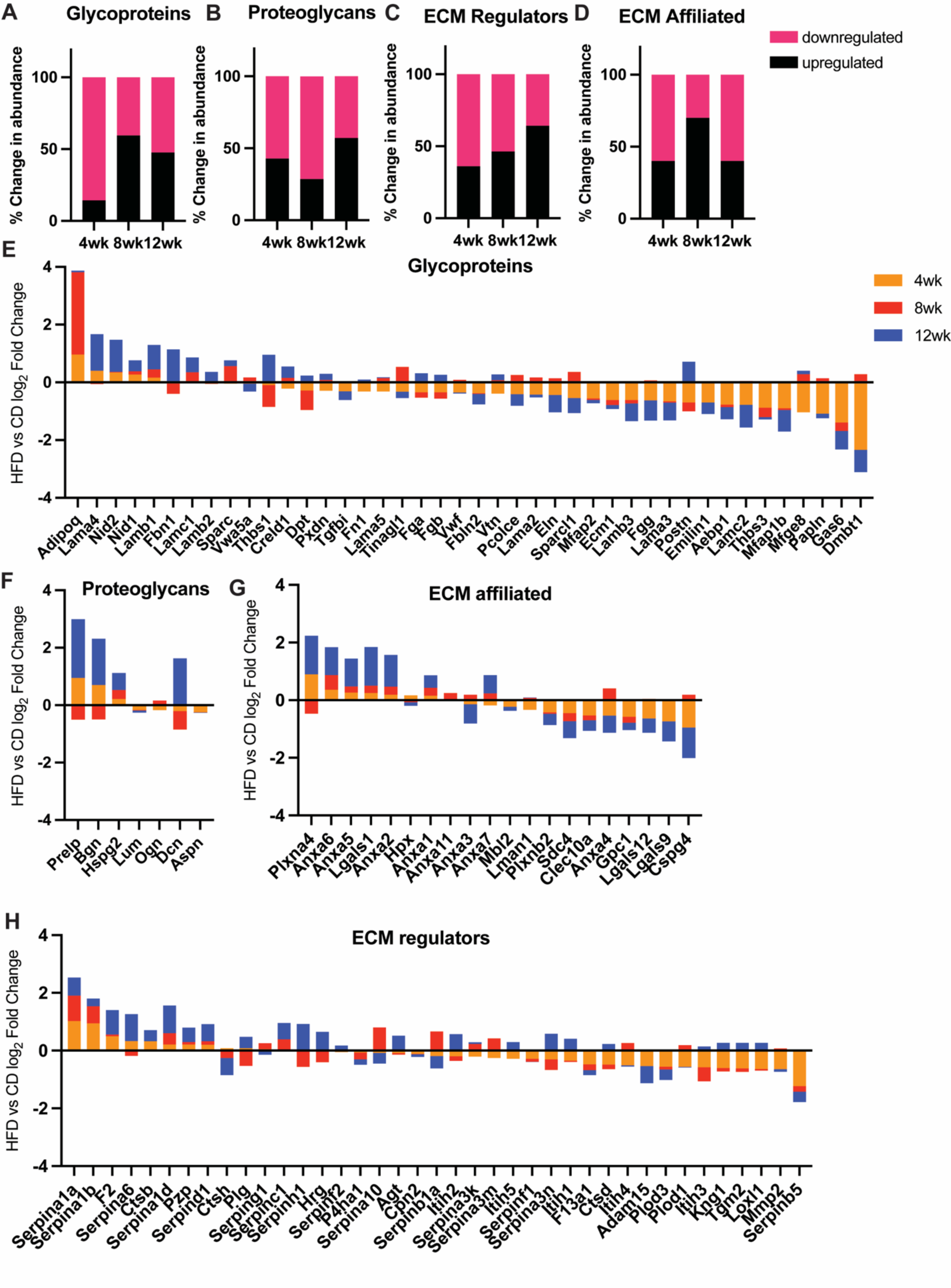
Relative abundance of glycoproteins, proteoglycans, ECM regulators, and ECM-affiliated proteins in the mammary fat pad on HFD. Percent change in abundance of upregulated and downregulated ECM proteins on HFD compared to age-matched CD **(A)** glycoproteins, **(B)** proteoglycans, **(C)** ECM regulators, and **(D)** ECM-affiliated. log_2_ Fold Change in ECM protein abundance in mammary fat pads on HFD compared to age-matched CD **(E)** glycoproteins, **(F)** proteoglycans, **(G)** ECM regulators, and **(H)** ECM-affiliated.

**Supplemental Figure 6.**
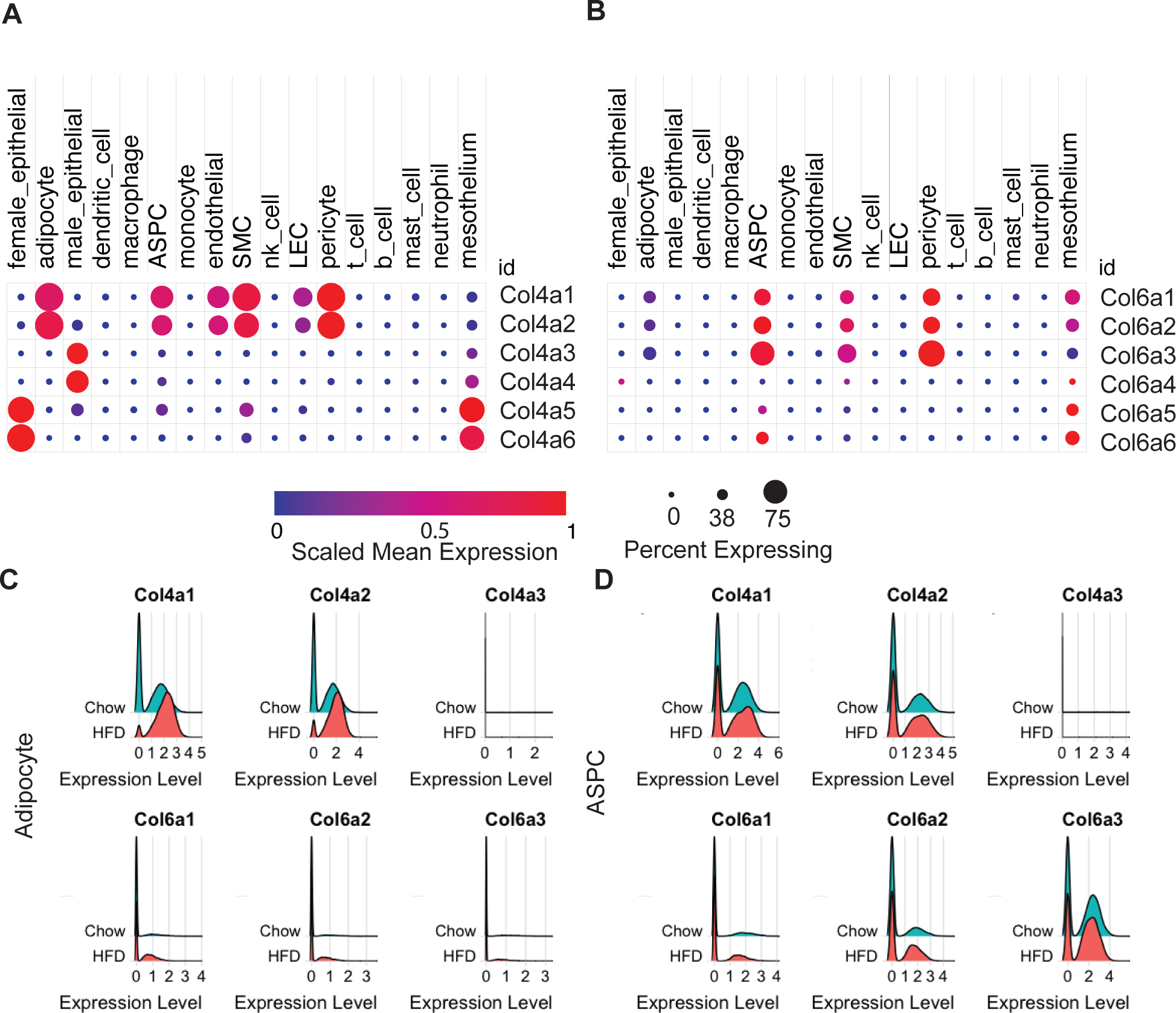
Mouse RNA-Seq data from mouse white adipose tissue. Scaled mean expression (blue low red high) and percentage of cells (size of dot) of murine white adipose tissue expressing **A)** Col6a1-6 and **B)** Col4a1-6. **C)** Col4a1-3 and Col6a1-3 expression by female murine subcutaneous adipocytes. **D)** Col4a1-3 and Col6a1-3 expression by female murine subcutaneous ASPC. Data from the Broad Institute [48].

**Supplemental Figure 7.**
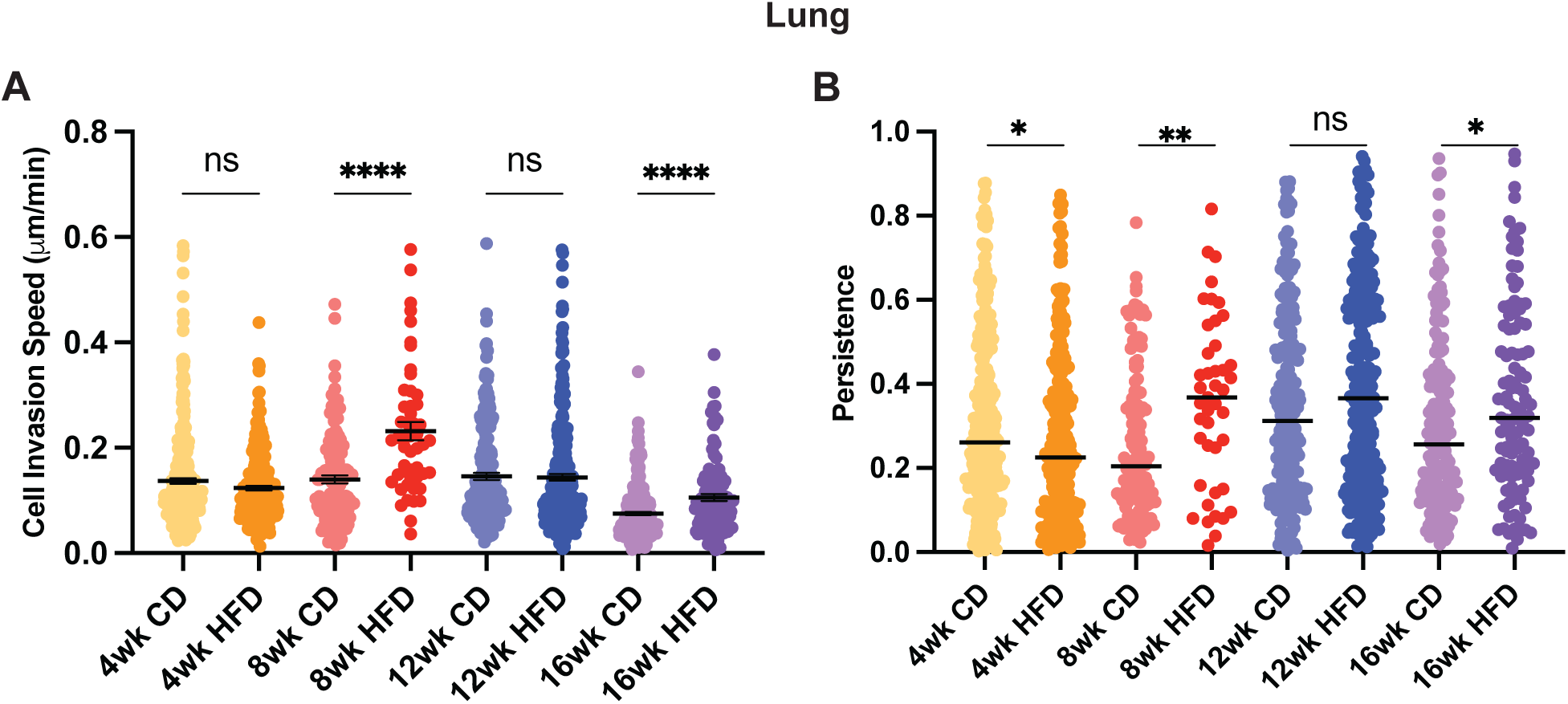
Age-matched CD and HFD Cell Invasion on lung ECM. **(A)** Cell invasion speed and **(B)** persistence of MDA-MB-231 cells invading on decellularized lung ECM from age-matched CD and HFD-fed mice for 4, 8, 12, and 16 weeks on diet. An unpaired t-test was used to compare age-matched CD and HFD groups (P*<0.05, P**<0.01, P***<0.001, P****<0.0001)

**Supplemental Figure 8.**
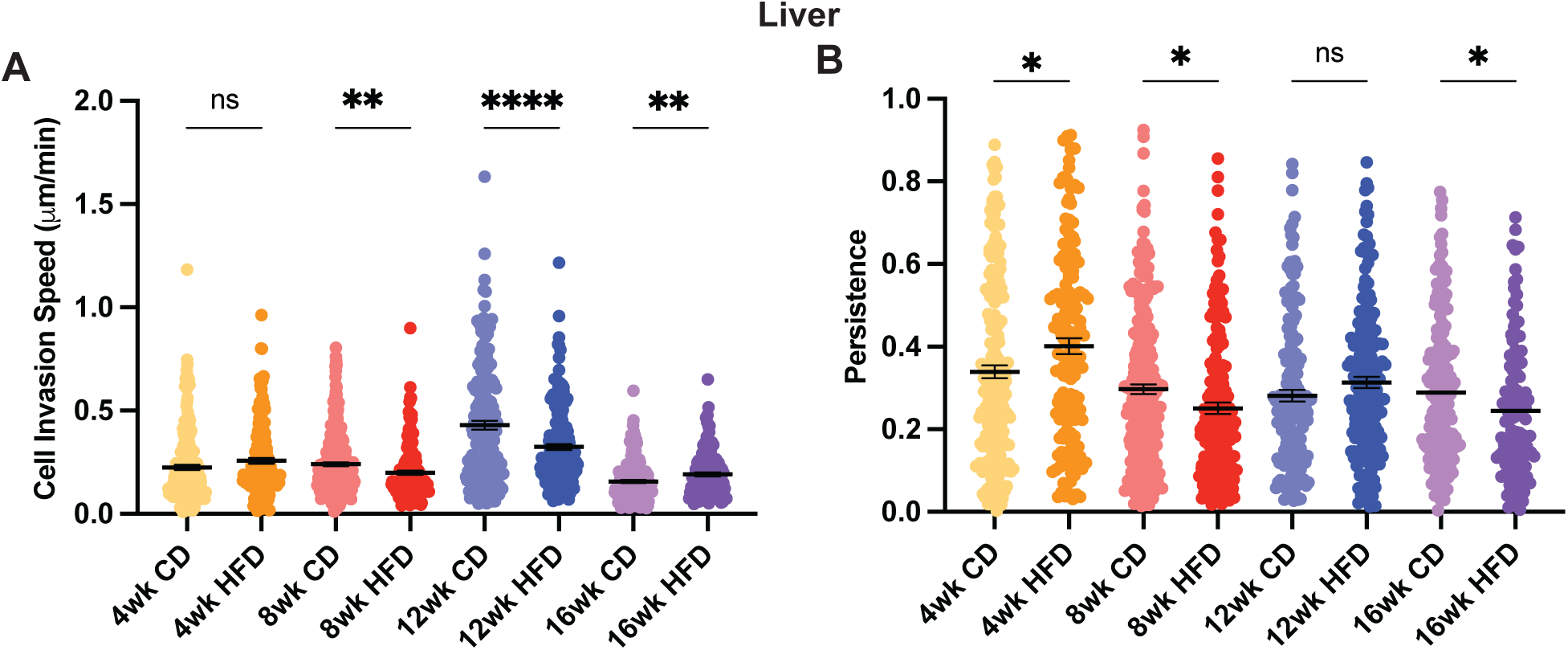
Age-matched CD and HFD Cell Invasion on liver ECM. **(A)** Cell invasion speed and **(B)** persistence of MDA-MB-231 cells invading on decellularized liver ECM from age-matched CD and HFD-fed mice for 4, 8, 12, and 16 weeks on diet. An unpaired t-test was used to compare age-matched CD and HFD groups (P*<0.05, P**<0.01, P***<0.001, P****<0.0001)

**Supplemental Figure 9.**
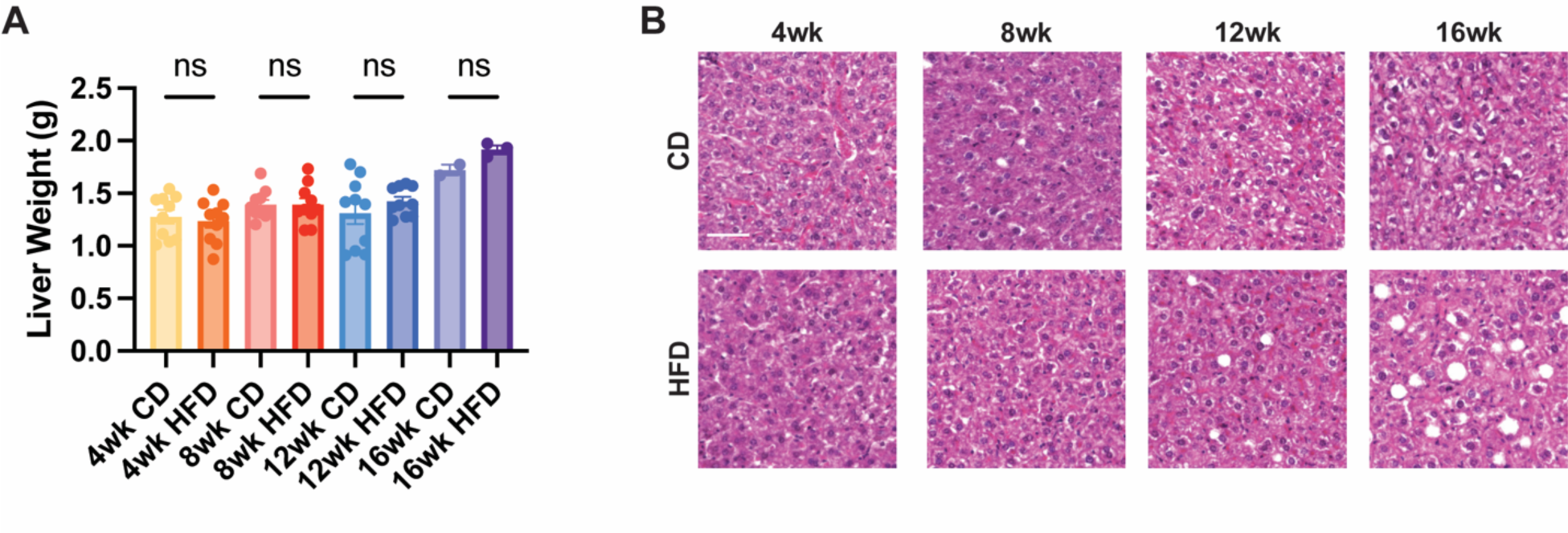
Liver weight and steatosis increase at later time points. **(A)** Liver weight (g) (**B)** H&E staining of liver tissue sections from mice fed HFD or CD for 4, 8, 12, and 16 weeks.

**Supplemental Figure 10.**
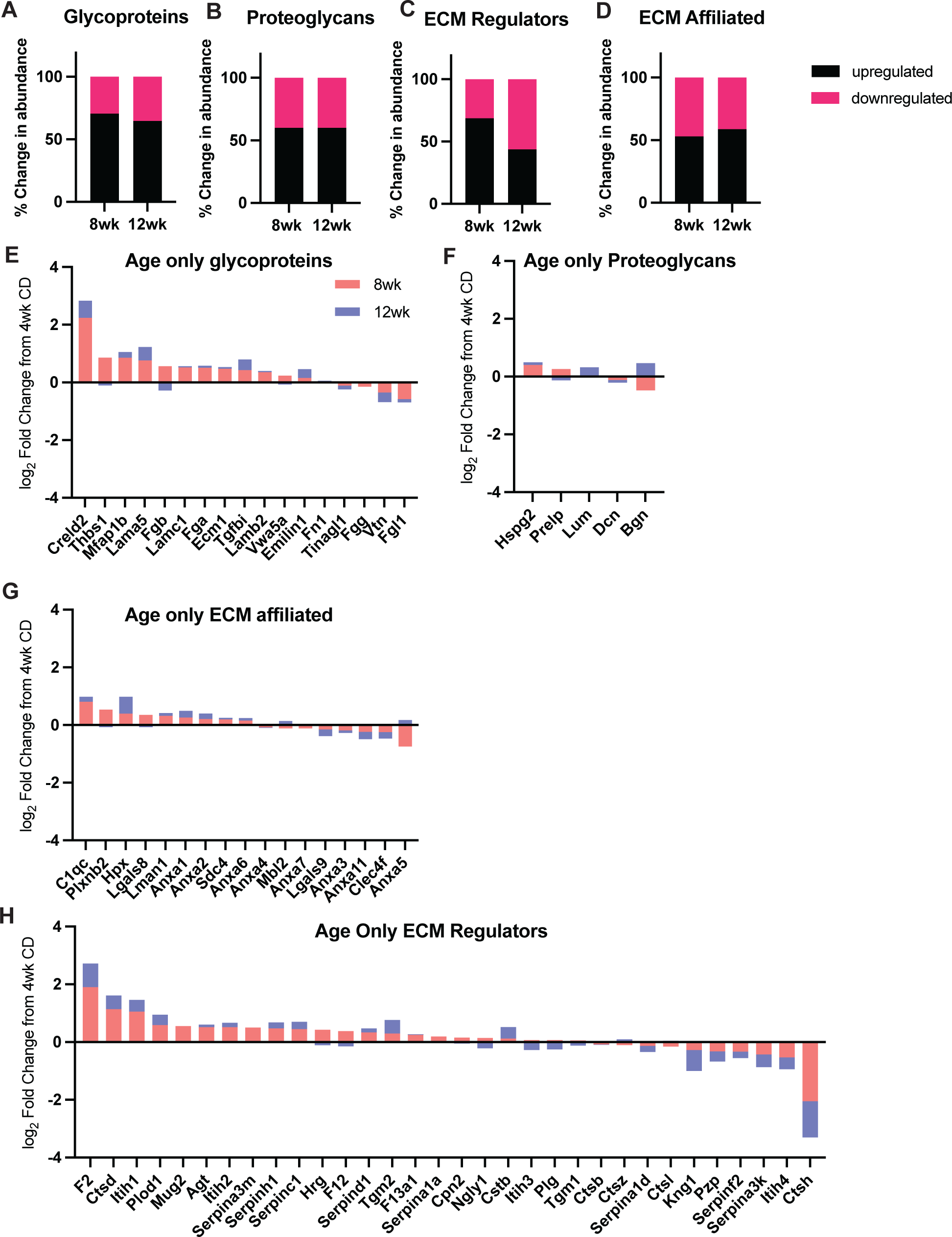
Relative abundance of glycoproteins, proteoglycans, ECM regulators, and ECM-affiliated proteins in the liver with age. Percent change in abundance of upregulated and downregulated ECM proteins with age compared to 4 weeks on CD **(A)** glycoproteins, **(B)** proteoglycans, **(C)** ECM regulators, and **(D)** ECM-affiliated. log_2_ Fold Change in ECM protein abundance in the liver with age compared to 4 weeks on CD **(E)** glycoproteins, **(F)** proteoglycans, **(G)** ECM regulators, and **(H)** ECM-affiliated.

**Supplemental Figure 11.**
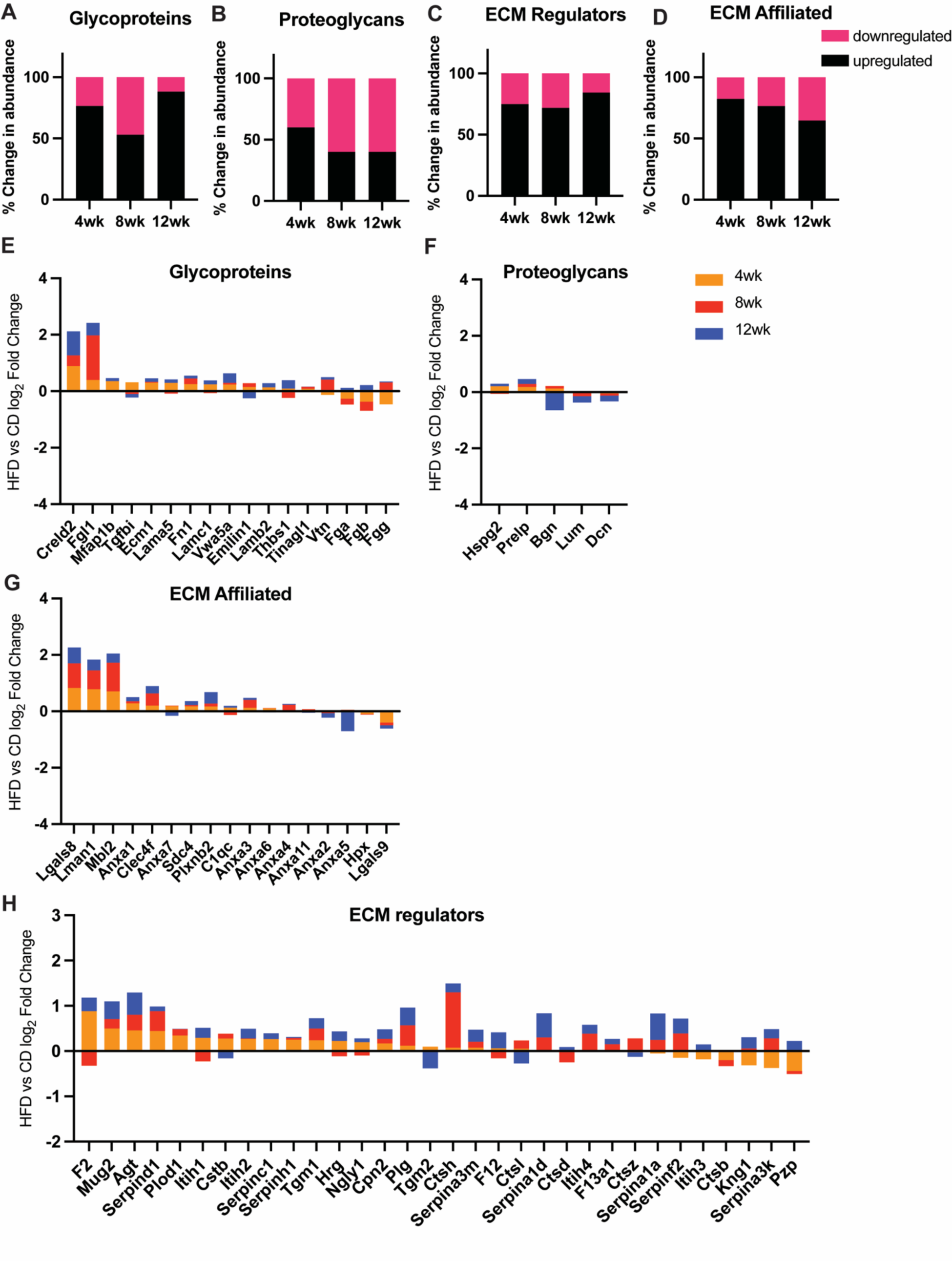
Relative abundance of glycoproteins, proteoglycans, ECM regulators, and ECM-affiliated proteins in the liver on HFD. Percent change in abundance of upregulated and downregulated ECM proteins on HFD compared to age-matched control **(A)** glycoproteins, **(B)** proteoglycans, **(C)** ECM regulators, and **(D)** ECM-affiliated. log_2_ Fold Change in ECM protein abundance in liver on HFD compared to age-matched control**(E)** glycoproteins, **(F)** proteoglycans, **(G)** ECM regulators, and **(H)** ECM-affiliated.

